# Ion-channel regulation of response decorrelation in a multi-scale model of the dentate gyrus

**DOI:** 10.1101/720078

**Authors:** Poonam Mishra, Rishikesh Narayanan

**Affiliations:** Cellular Neurophysiology Laboratory, Molecular Biophysics Unit, Indian Institute of Science, Bangalore 560012, India.

**Keywords:** channel decorrelation, computational model, hippocampus, intrinsic plasticity, ion channels

## Abstract

The dentate gyrus (DG) is uniquely endowed with multiple forms of biological heterogeneities owing to the expression of adult neurogenesis and sparse connectivity, and has been functionally implicated in response decorrelation and pattern separation. Although channel decorrelation could be achieved through synergistic interactions between these heterogeneities, the impact of individual ion channels on channel decorrelation has not been explored. Here, to systematically assess the cascading impact of molecular-scale (ion channel) perturbations on network-scale outcomes (decorrelation), we first quantified the impact of eliminating individual ion channels on single-cell physiology of heterogeneous populations of granule cells (GCs) and basket cells (BCs). Employing virtual knockout simulations involving both populations, we found that the mapping between ion channels and nine distinct physiological measurements was many-to-many. Next, to assess the impact of ion channel elimination on channel decorrelation, we employed a conductance-based multi-scale network model of the DG. This network was endowed with four distinct forms of heterogeneities (intrinsic, synaptic, structural and afferent), with afferent inputs from the entorhinal cortices driven by virtual arena traversal. We show that individual ion channels expressed in GCs govern DG network excitability, and critically regulate the ability of the network to perform channel decorrelation. The impact of eliminating individual ion channels on network excitability and channel decorrelation was differential and variable, with local heterogeneities playing a pivotal role in determining the strength of such impact. Specifically, in the presence of structurally immature neurons in the DG network, the impact of ion channel elimination on channel decorrelation was considerably lower when compared with a network exclusively constructed with structurally mature neurons. Finally, we show that for any given ion channel knockout, the average percentage change in output correlation was invariant to the specific values of input correlation, across different network configurations endowed with disparate structural and afferent heterogeneities. Our analyses emphasizes that the mapping between components and function is many-to-many across scales, and assign critical roles for biological heterogeneities in conferring multi-scale functional robustness in the face of physiological and pathological perturbations.

**SIGNIFICANCE STATEMENT:** There are precise sets of computation spanning different scales of analyses that drive behavioral states and responses of an animal. Perturbations to components that drive these computations at one scale could result in a cascading effect that alters physiological properties across several scales. Multi-scale computational models that account for biological heterogeneities at each scale are ideal tools to approach this complex problem, where systematic analyses of such cascades could be rigorously accomplished. Here, we systematically assessed the impact of eliminating individual ion channels, first on neuronal intrinsic properties, and consequently on network excitability and response decorrelation. Our results unveil important roles for biological heterogeneities in conferring multi-scale functional robustness in the face of physiological and pathological perturbations, achieved through many-to-many mappings between constitutive components and physiological outcomes.

## INTRODUCTION

There are precise sets of computation spanning different scales of analyses that drive behavioral states and responses of an animal. Perturbations to components that drive these computations at one scale, introduced by pathological insults or neuromodulation or learning- or adaptation-induced changes, could result in a cascading effect that alters physiological properties across several scales. Experimental analyses of such cascades employing either pharmacological agents or genetic manipulations of components is complicated by the presence of non-specificities in the pharmacological blockade or compensations consequent to the elimination of these ion channels. Therefore, computational models spanning different scales, where a systematic analyses of such cascades can be rigorously accomplished, provides an ideal path to approach this problem. In this study, we systematically assessed the cascading impact of eliminating individual ion channels from two distinct neuronal subtypes, first on neuronal intrinsic physiological properties, and consequently on network excitability and on the ability of the network to perform response decorrelation.

The dentate gyrus (DG) is uniquely endowed with multiple forms of biological heterogeneities, owing to the expression of adult neurogenesis and the sparse, orthogonal and divergent nature of connectivity. Biological heterogeneities that span the DG include those in ion channel properties and expression profiles, neuronal intrinsic properties, dendritic arborization, local synaptic connectivity and afferent connectivity. Importantly, each of these heterogeneities is amplified by the expression of adult neurogenesis, with the enormous dependence of each of these individual properties on the neuronal maturation process. Uniquely, with reference to afferent connectivity, adult neurogenesis provides a substrate for orthogonal, non-overlapping processing and storage of afferent information from upstream cortex. The expression of these amplified heterogeneities, in conjunction with specific characteristics of the local network has resulted in postulates and lines of evidence that the DG network is an ideal system to execute channel decorrelation and pattern separation (Aimone et al., 2006; Leutgeb et al., 2007; Aimone et al., 2009; Sahay et al., 2011; Dieni et al., 2013; Aimone et al., 2014; Kropff et al., 2015; Li et al., 2017; Lodge and Bischofberger, 2019; Mishra and Narayanan, 2019).

Although it has been established that channel decorrelation could be achieved through synergistic interactions between different forms of these heterogeneities (Mishra and Narayanan, 2019), the impact of individual ion channels on channel decorrelation or their interactions with these heterogeneities has not been studied. A dominance hierarchy has been established with reference to impact of different heterogeneities on channel decorrelation, with afferent heterogeneities dominating local heterogeneities when they are coexpressed (Mishra and Narayanan, 2019). Against this backdrop, if there were ion channel perturbations to neurons in the local network, would the expression of different forms of heterogeneities contribute to functional resilience of the network? We employed a multi-scale conductance-based network model of the DG, endowed with four distinct forms of local and afferent heterogeneities, and systematically assessed the cascading impact of eliminating individual ion channels on neuronal and network physiology.

At the single-neuron scale, our analyses revealed that the mapping between ion channels and physiological measurements was many-to-many. At the network scale, the impact of eliminating individual ion channels was differential and variable both in terms of affecting network firing rates and channel decorrelation, and was critically reliant on the specific local heterogeneities expressed in the DG network. Importantly, in the presence of structurally immature neurons in the DG network, the impact of ion channel elimination on channel decorrelation was considerably lower when compared with a network constructed with structurally mature neurons. Finally, we showed that for a given ion channel, the average percentage change in output correlation was invariant to the specific values of input correlation, across different network configurations endowed with disparate structural and afferent heterogeneities. Our analyses emphasizes that the mapping between components and function is many-to-many across scales, and assign critical roles for heterogeneities in conferring multi-scale functional robustness in the face of physiological and pathological perturbations.

## METHODS

In this multi-scale computational study, we sought to systematically examine the impact of different ion channel subtypes on neuronal intrinsic properties and on network-scale decorrelation in neuronal responses in the DG. In what follows, we describe the models, procedures and measurements employed in assessing single-neuron and network physiology after virtual knockout of specific ion channels. To avoid confusion between the use of the word “channel” in channel decorrelation and ion channels, we always employ the phrase “ion channels” when we refer to the latter.

### Impact of virtual knockout of different ion channels on single-neuron intrinsic properties: Assessment in a heterogeneous population of granule and basket cell models

Analyzing the impact of individual ion channels on neuronal intrinsic properties in a single hand-tuned model would introduce biases that are inherent to the specific model and would not account for the heterogeneities in ion channel expression or in intrinsic properties of the neurons. With the ubiquitous expression of ion-channel degeneracy, whereby synergistic interactions among disparate combinations of channels result in the emergence of similar single neuron physiological characteristics, such an approach would yield results that are not applicable to the entire population of neurons in the biological system. A well-established alternate to this approach, which accounts for degeneracy and heterogeneities across scales, is an unbiased stochastic search algorithm that spans the ion channel parametric space to arrive at neuronal models that satisfy cellular-scale physiological constraints. As the population of neuronal models arrived through such an approach is well constrained by the biophysical and physiological measurements from the specific neuronal subtype under consideration, this population has been employed to understand the impact of individual ion channels on neuronal physiology. An important difference between a hand-tuned model and this stochastic search approach is that the stochastic search model is unbiased, with no relationship in terms of *which* channel was introduced for matching *what* specific physiological property. Therefore, results arrived on the impact of individual channels *emerge* from the heterogeneous neuronal populations, without assignment of specific physiological purposes to individual channels (Foster et al., 1993; Goldman et al., 2001; Prinz et al., 2003; Taylor et al., 2009; Marder and Taylor, 2011; Rathour and Narayanan, 2012a, 2014; Mishra and Narayanan, 2015; Srikanth and Narayanan, 2015; Basak and Narayanan, 2018; Mittal and Narayanan, 2018; Rathour and Narayanan, 2019).

Therefore, we employed the multi-parametric multi-objective stochastic search (MPMOSS) as an ideal route to generate a heterogeneous population of GC and BC neuronal models. In this study, to assess the impact of individual ion channels on neuronal and network physiology, we employed the valid models generated in our previous study (Mishra and Narayanan, 2019). We briefly describe the details associated with the generation of these heterogeneous populations of single compartmental neuronal models. The stochastic search for valid granule cells involved 40 active parameters associated with passive properties, nine active conductances and calcium handling mechanisms. The nine different active conductances that were present in our GC model were (Fig. 1*A*): hyperpolarization-activated cyclic nucleotide gated (HCN or *h*), *A*-type potassium (KA), fast sodium (NaF), delayed-rectifier potassium (KDR), small conductance (SK) and big conductance calcium-activated potassium (BK), *L*-type calcium (CaL), *N*-type calcium (CaN) and *T*-type calcium (CaT). The channel kinetics and their voltage-dependent properties were adopted from experimental measurements from the GC (Beck et al., 1992; Magee, 1998; Aradi and Holmes, 1999; Ferrante et al., 2009). The evolution of cytosolic calcium concentration [*Ca*]*_c_*, was dependent on the current through voltage-gated calcium channels and involved a first order decay with a default calcium decay time constant, τ_Ca_=160 ms. In generating the physiologically-validated heterogeneous GC population, we subjected 20,000 unique models spanning this 40-parameter space to a validation procedure involving nine different single-cell electrophysiological measurements (see below) from GCs. We found 126 models (∼0.63% of the total population) to be valid.

**Figure 1.**
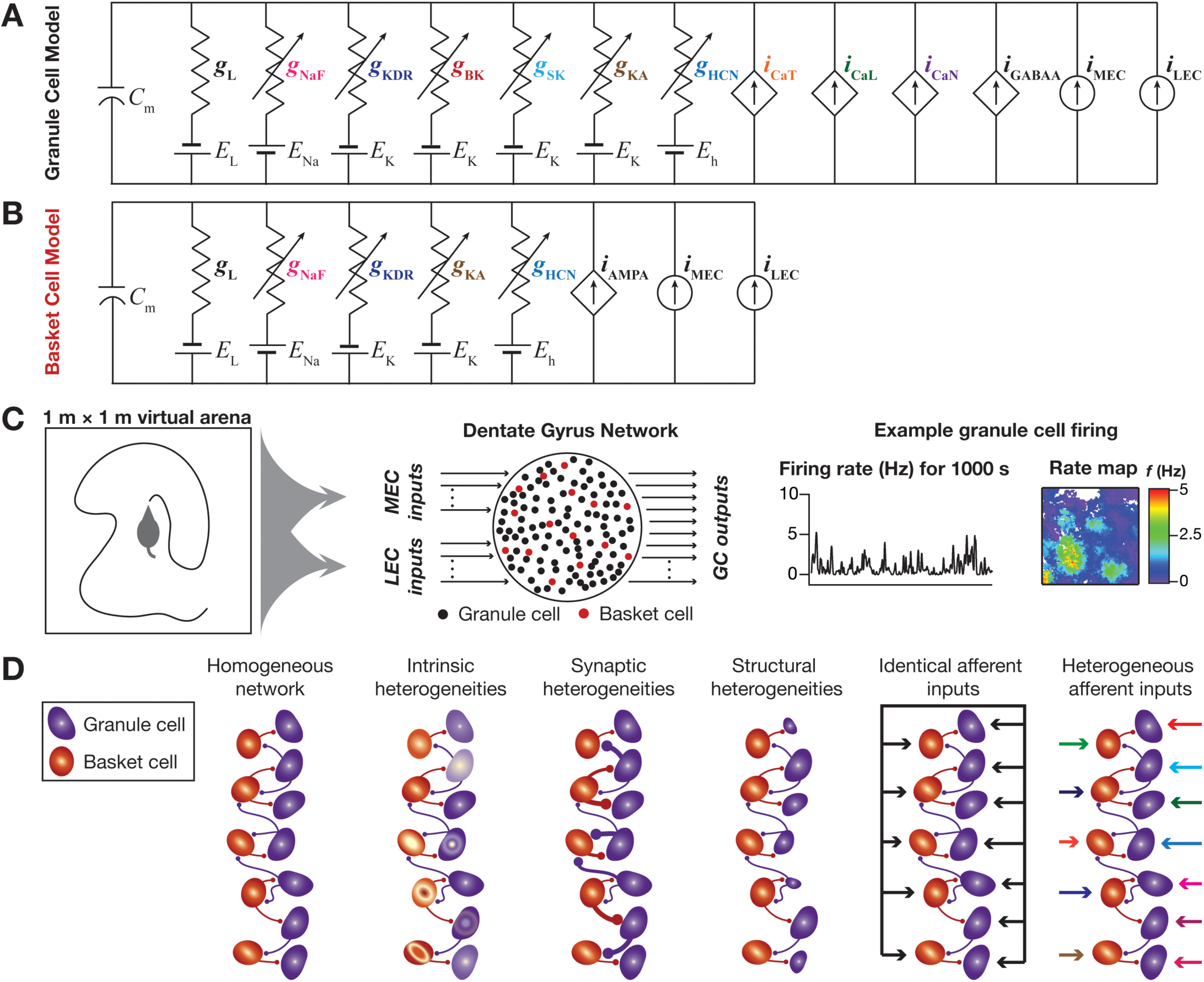
Multi-scale modeling framework for assessing the impact of ion channel knockouts on cellular and network physiology of the dentate gyrus during virtual arena traversal. *A–B*, Conductance-based models for granule cells (GC) and basket cells (BC) were built using several experimentally derived ion channel conductances for these neurons. Symbols employed: *g*_L_: Leak conductance; *g*_NaF_: Fast sodium conductance; *g*_KDR_: Delayed rectifier potassium conductance; *g*_BK_: Big-conductance calcium-activated potassium conductance; *g*_SK_: Small-conductance calcium-activated potassium conductance; *g*_BK_: *A*-type transient potassium conductance; *g*_HCN_: Hyperpolarization-activated cyclic-nucleotide-gated (HCN) nonspecific cation conductance; *i*_CaT_: *T-*type calcium current; *i*_CaL_: *L-*type calcium current; *i*_CaN_: *N-*type calcium current; *i*_GABAA_: GABA_A_ receptor current; *i*_AMPA_: AMPA receptor current; *i*_MEC_: current from medial entorhinal cortical cells; *i*_LEC_: current from lateral entorhinal cortical cells. Note that all sodium, potassium and nonspecific cation channels were modeled using a Nernstian framework are represented as parallel conductances; all calcium currents and receptor currents were modeled using the Goldman-Hodgkin-Katz (GHK) formulation, and therefore are represented as dependent currents; the inputs from entorhinal cortex were modeled as currents that were dependent on animal traversal and are represented as current sources. *C*, A virtual animal was allowed to run in an arena of 1 m × 1 m (left panel) for a period of 1000 s to allow complete traversal of the entire arena. The animal’s location in this arena was fed into a dentate gyrus network made of interconnected granule and basket cells (middle panel). The voltage outputs of granule cells were recorded to obtain firing rate profiles and spatial rate maps (last panel) by overlaying neuronal firing rate over the temporally aligned spatial location of the virtual animal. *D,* The network employed here was not a homogeneous network, but employed several biological heterogeneities expressed in the dentate gyrus. Intrinsic heterogeneities represented variability in ion channel densities and neuronal intrinsic properties, and was accounted for by employing a multi-parametric multi-objective stochastic search (MPMOSS) paradigm (Mishra and Narayanan, 2019). Synaptic heterogeneities represented the strength of the local BC → GC and GC → BC connections, and were modeled by altering the AMPAR and GABA_A_ receptor permeability. These receptor permeabilities were varied within a range where the excitation-inhibition balance was maintained and the overall firing rates of GCs and BCs were within experimentally observed ranges. Structural heterogeneities were introduced to model surface area changes in granule cells consequent to adult neurogenesis, and were incorporated by adjusting the geometry of the GC models. Afferent heterogeneities were representative of the uniquely sparse connectivity from the entorhinal cortices to the DG, and were modeled by feeding each GC and BC neuron with different afferent inputs. This scenario was compared with a case where all GCs and BCs were given *identical* afferent inputs.

A similar MPMOSS strategy was employed to generate a heterogeneous population of basket cells (Fig. 1*B*), endowed with four different voltage-gated ion channels (HCN, KA, NaF and KDR), and involving a stochastic search space of 18 parameters (ion channel and passive membrane properties). Here, we generated 8,000 unique BC models, validated them against 9 electrophysiological measurements from BCs and found 54 valid BC models (∼0.675% of the total population). The experimental bounds on measurements for granule and basket cells were obtained from (Mott et al., 1997; Lubke et al., 1998; Aradi and Holmes, 1999; Santhakumar et al., 2005; Krueppel et al., 2011). We have demonstrated ion channel degeneracy individually for the 126 valid GCs and for the 54 BCs (Mishra and Narayanan, 2019), whereby the underlying parametric combinations that yielded these model populations exhibited tremendous variability across models. We employ these valid model populations for the virtual knockout analyses presented in this study.

### Single-neuron intrinsic properties and measuring the impact of virtual ion-channel knockouts on neuronal physiology

The intrinsic response properties of GCs and BCs were quantified based on nine measurements (Lubke et al., 1998; Mishra and Narayanan, 2019), which were employed to validate the models obtained through stochastic search and to assess the impact of individual channel knockouts. The nine measurements, which were computed employing procedures identical to those described in the literature (Mishra and Narayanan, 2019), are neuronal firing rate with pulse-current injections of 50 pA (*f*_50_) or 150 pA (*f*_150_), sag ratio, input resistance (*R*_in_), action potential (AP) amplitude (*V*_AP_), AP threshold (*V*_th_), AP half-width (*T*_APHW_), fast afterhyperpolarization (*V*_fAHP_) and spike frequency adaptation (SFA). *R*_in_ was measured from the neuronal steady state voltage response to each of 11 different current pulses, injected with amplitudes ranging from –50 pA to 50 pA (for 1000 ms) in steps of 10 pA (*e.g.*, Fig. 2*A*, left). The steady state voltage deflections from *V*_RMP_ were plotted as a function of the corresponding current injections to obtain a *V–I* plot. We fitted a straight-line function to this *V–I* plot, and the slope of this linear fit defined *R*_in_. Sag ratio was calculated as the ratio of the steady state voltage deflection to the peak voltage deflection recorded in response to a –50 pA (1000 ms) current injection (Mishra and Narayanan, 2019).

**Figure 2.**
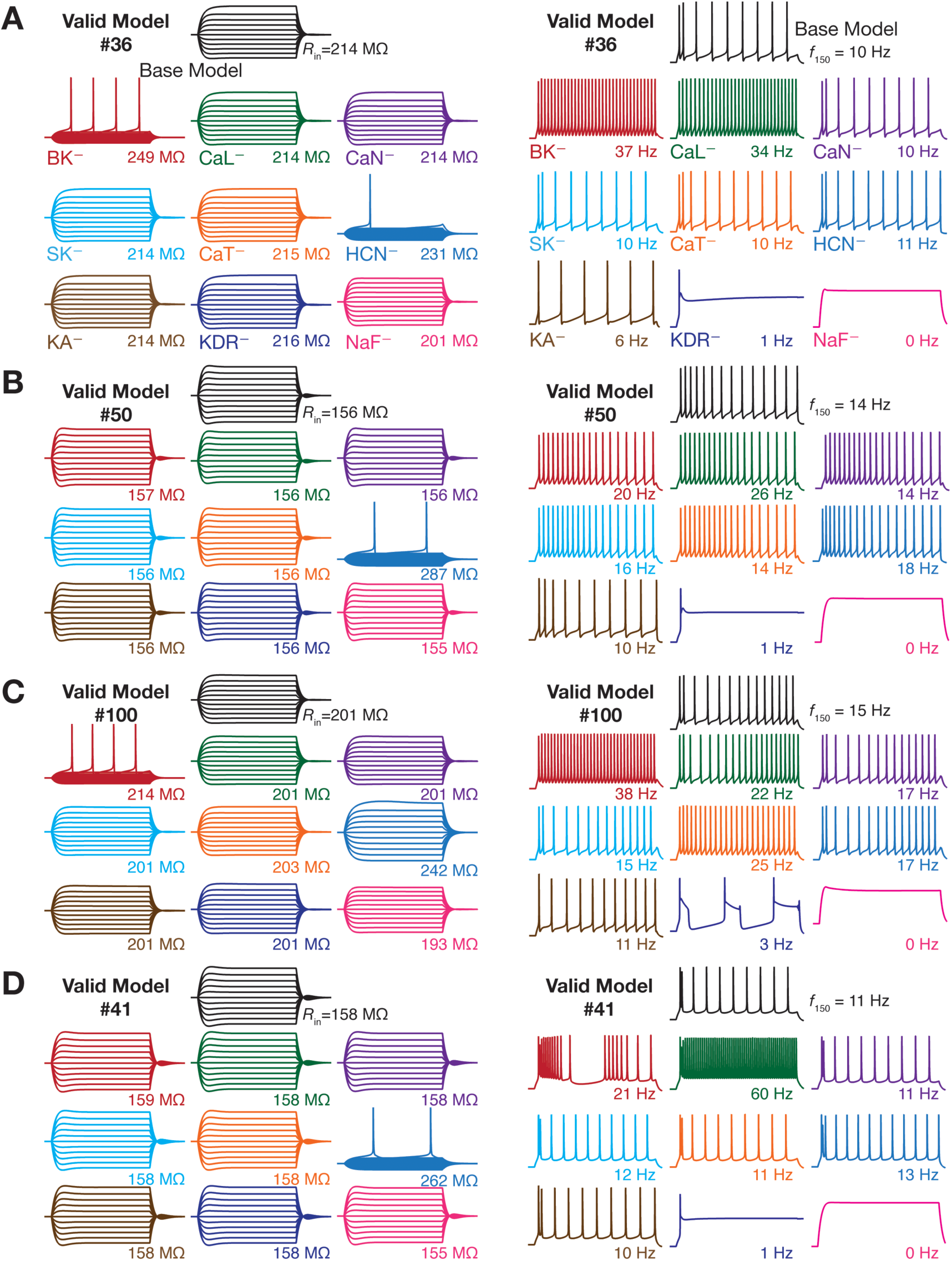
Virtual knockout of individual channels resulted in heterogeneous and differential impact on different sub- and supra-threshold electrophysiological measurements in a valid population of dentate granule cells. *A*, *Left*: Voltage responses to current pulses of –50 to +50 pA, in steps of 10 pA, for 500 ms employed for input resistance (*R*_in_) calculation. *Right:* Voltage traces showing firing rate (*f*_150_) and spike pattern in response to a 150 pA current injection for 950 ms for the same granule cell model. *B–D*, Same as (*A*) but for different valid models of granule cell. Valid models 36 (*A*) and 50 (*B*), respectively represent the minimum and maximum percentage change in input resistance (see Fig. 3G) after virtual knockout of HCN ion channels. Valid models 100 (*C*) and 41 (*D*), respectively represent the minimum and maximum percentage change in *f*_150_ (see Fig 3F) after virtual knockout of *L-*type calcium ion channels. Across panels, black traces represent the valid base model and traces of other colors depict those after virtual knockout of nine different ion channels in the chosen model.

All supra-threshold measurements were obtained from the voltage trace recorded in response to a 150 pA depolarizing current injection, with AP measurements obtained from the first spike of this trace. Firing frequency was calculated as number of spikes in response to 150 pA current injection for one second (*e.g.*, Fig. 2*A*, right). Spike frequency adaptation (SFA) was calculated as the ratio of the first inter spike interval (ISI) to the last ISI. The voltage in the AP trace corresponding to the time point at which the d*V*/d*t* crossed 20 V/s defined AP threshold. AP half-width was the temporal width measured at the half-maximal points of the AP peak with reference to AP threshold. AP amplitude was computed as the peak voltage of the spike relative to *V*_RMP_. Fast afterhyperpolarization (*V*_fAHP_) was measured as the maximal repolarizing voltage deflection of the AP from threshold (Mishra and Narayanan, 2019).

As disparate parametric combinations yielded similar physiological properties in our heterogeneous population of GC and BC models (Mishra and Narayanan, 2019), it was important to independently assess the impact of channel elimination in each of the 126 GCs and 54 BCs. Within the degeneracy framework, virtual knockout models (VKMs) constitute a powerful technique to quantitatively assess the contribution of specific channels to chosen measurements in a heterogeneous population of models (Rathour and Narayanan, 2014; Anirudhan and Narayanan, 2015; Mukunda and Narayanan, 2017; Basak and Narayanan, 2018). Specifically, for the GC population, we virtually knocked-out one of the 9 active channels (by setting its conductance value to be zero) individually from each of the 126 valid models, and computed each of the 9 measurements after this knockout. Then, we computed the percentage change in each of 9 measurements from their respective valid base model values (when all the channels were intact in that specific valid model). This procedure was repeated for all nine channels in GCs, and the statistics of percentage changes in each measurement for each VKM were assessed. A similar procedure was applied independently on BC valid models as well, with differences in number of active channels in the model (4) and number of valid models (54). The procedure was employed to assess each of the 9 measurements here as well.

Quantitatively, for each of the 9 different measurements, let *M*_n_ (*base*) represent the measurement value of the base version (*i.e.*, all parameters were intact) of model number *n* (1 ≤ *n* ≤ 126 for GCs; 1 ≤ *n* ≤ 54 for BCs). Let *M*_n_ (*C*_i_) represent the measurement value obtained from the VKM *after* deleting one the channels *C*_i_ (1 ≤ *i* ≤ 9 for GCs; 1 ≤ *i* ≤ 4 for BCs). We quantified the impact of single ion channel knockout on each measurement by computing percentage change as:

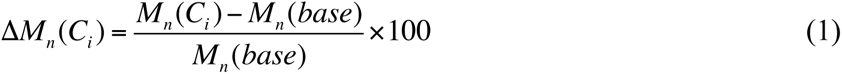

This procedure was repeated for each valid model, each channel and each measurement, and the statistics of these measurements were plotted as quartiles to depict the entire span of changes (Fig. 3).

**Figure 3.**
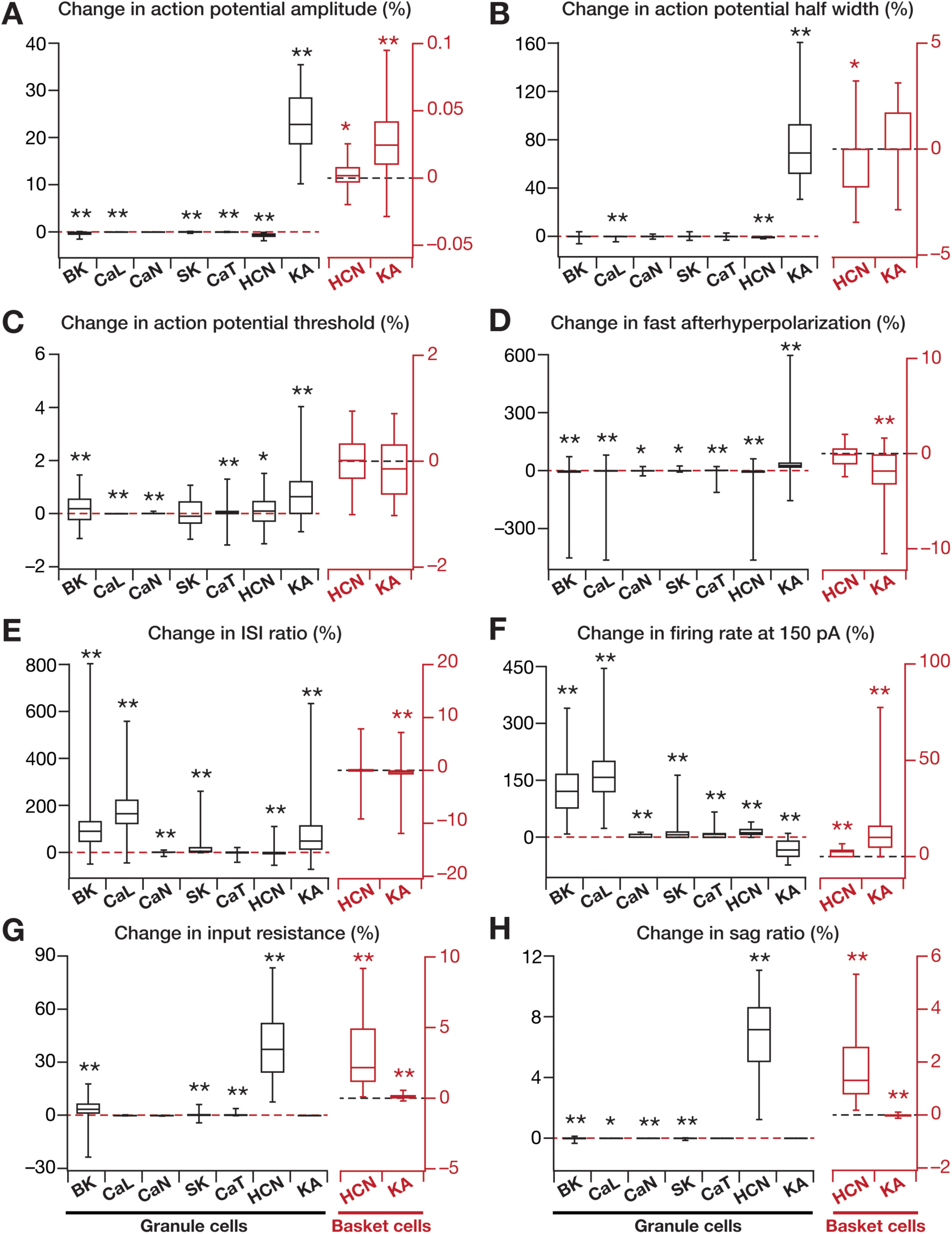
The mapping between individual ion channels and different electrophysiological measurements was many-to-many, with virtual knockout of individual ion channels yielding differential and variable effects on different measurements. *A–H*, Plots of percentage changes in different electrophysiological measurements obtained after virtual knockout of individual ion channels from valid models of granule (*N*_valid_ = 126 for GC, black) and basket (*N*_valid_ = 54 for GC, red) cell population obtained using MPMOSS. Percentage changes were calculated by comparing the measurement after virtual knockout of the specific ion channel with the measurement in the corresponding base model. Individual panels represent the following intrinsic measurements: *A,* action potential amplitude, *V*_AP_; *B,* action potential half-width, *T*_APHW_; *C,* action potential threshold, *V*_th_; *D,* fast after hyperpolarization potential, *V*_AHP_; *E*, ISI ratio; *F,* firing rate in response to 150 pA current injection, *f*_150_; *G,* input resistance, *R*_in_; and *G,* sag ratio. *p* values were obtained using Wilcoxon signed rank test, where the percentage change in measurements were tested for significance from a “no change” scenario. *: *p* < 0.01, **: *p* < 0.001.

### The multi-scale network and its inputs

A network of 100 GCs and 15 BCs (Fig. 1*C*), with the GC:BC ratio constrained by experimental observations (Aimone et al., 2009), was constructed by randomly picking valid models from the population of GCs and BCs obtained from MPMOSS (Mishra and Narayanan, 2019). Local connectivity was set such that the probability of a BC to GC connection was 0.1, and that of a GC to BC connection was set as 0.05 (Aimone et al., 2009). The GC → BC and BC → GC connections were modeled as synapses containing AMPA and GABA_A_ receptors, respectively. The AMPA receptors were modeled to be permeable to sodium and potassium ions, whereas the GABA_A_ receptors were permeable to chloride ions. Both receptor currents were modeled using the GHK convention (Goldman, 1943; Hodgkin and Katz, 1949), with a rise time of 2 ms and a decay time constant of 10 ms (Mishra and Narayanan, 2015, 2019).

All neurons in this DG network received inputs from two different regions of entorhinal cortex (EC): one from medial entorhinal cortex (MEC) grid cells that transmitted spatial information and another from lateral entorhinal cortex (LEC), which provides contextual information (Anderson et al., 2007; Renno-Costa et al., 2010). The firing of the MEC and LEC cells were driven by the position of a randomly traversing virtual animal in a 1 m × 1 m arena (Fig. 1*C*). Each DG neuron received *active* inputs (Cayco-Gajic et al., 2017; Mishra and Narayanan, 2019) from 5 different MEC cells and 5 different LEC cells, with input strength from MEC and LEC split equally. The current input from a single MEC grid cell to DG cells was modeled as a hexagonal grid function defined as a sum of three two-dimensional cosine functions (Solstad et al., 2006). This hexagonal grid function was scaled to set the relative contribution of MEC and LEC to DG cells. Each MEC cell input was distinct in terms of the grid frequency and grid orientation, each randomly sampled from respective uniform distributions. For modeling LEC inputs to GCs and BCs, we tiled the arena into 25 squares (5 rows and 5 columns) and assigned different tiles to be active for different LEC inputs (Renno-Costa et al., 2010). Inputs from this LEC cell to the DG cell was then scaled to set equal relative contribution of MEC and LEC to the DG cells. Each LEC cell input was associated with a unique randomized matrix, representing different active and inactive regions (Renno-Costa et al., 2010; Mishra and Narayanan, 2019).

### Incorporating different forms of heterogeneities into the multi-scale network

To assess the robustness of networks endowed with different forms of heterogeneities to single channel knockouts, we built DG networks endowed with distinct combinations of four different types of heterogeneities (Fig. 1*D*), following the approaches introduced in (Mishra and Narayanan, 2019):

i. *intrinsic heterogeneity*, where the GC and BC model neurons had widely variable intrinsic parametric combinations, yet yielded physiological measurements that matched their electrophysiological counterparts. As mentioned in earlier sections, this was incorporated into our network by the use of independent MPMOSS algorithms for BCs and GCs (Mishra and Narayanan, 2019).
ii. *synaptic heterogeneity*, where the synaptic strength of the local GC-BC network was variable with excitatory and inhibitory synaptic permeability values picked from uniform random distributions. These parameters were maintained at a regime where the peak-firing rate of GCs and BCs stayed within their experimental ranges of 4–10 Hz and 30–50 Hz, respectively (Leutgeb et al., 2007). We ensured that extreme parametric combinations where the cell ceased firing (because of depolarization-induced block at one extreme or high inhibition at the other) were avoided, implying a balance between excitatory and inhibitory connections (Mishra and Narayanan, 2019).
iii. *neurogenesis-induced structural heterogeneity*, where the DG network was constructed entirely of mature or immature neurons, or constructed from neurons that represented different randomized neuronal ages. Populations of immature GCs (originating through adult neurogenesis) were obtained by subjecting the mature set of the GC valid models (obtained through MPMOSS) to structural plasticity. Specifically, the reduction in dendritic arborization and in the overall number of channels expressed in immature neurons (Aimone et al., 2014) was approximated by a reduction in the surface area (diameter) of the model neuron, using *R*_in_ as the measurement to match with experimental counterparts. The diameters of GC for the three distinct configurations were: fully mature (63 µm), fully immature (BCs: 1–3 µm) and heterogeneous age population (2–63 µm). Neurogenesis-induced structural heterogeneity was confined to the GC population, leaving the BC population to be mature (Mishra and Narayanan, 2019).
iv. *input-driven or afferent heterogeneity*, where all neurons in the GC and BC populations received either *identical* inputs (absence of afferent heterogeneity) from the EC, or *each* GC and BC received unique inputs (presence of afferent heterogeneity) from the EC (Mishra and Narayanan, 2019). Afferent heterogeneity models incorporate sparse and orthogonal afferent connectivity from the EC to the DG (Aimone et al., 2006; Andersen et al., 2006; Amaral et al., 2007; Aimone et al., 2009; Aimone et al., 2010, 2011; Aimone and Gage, 2011; Aimone et al., 2014; Li et al., 2017; Lodge and Bischofberger, 2019).

We tested the impact of virtually knocking out individual ion channels in networks endowed with different combinations of biological heterogeneities, to ensure that our conclusions were not reflections of narrow parametric choices and to ask if the expression of heterogeneities enhances the robustness of the network to ion channel perturbations. There are several lines of evidence that the synaptic connectivity to immature neurons are low, and that this low connectivity counterbalances their high excitability (Mongiat et al., 2009; Dieni et al., 2013; Dieni et al., 2016; Li et al., 2017). To account for these, we reduced the overall afferent drive in scenarios that involved neurogenesis-induced structural differences (*i.e.*, the fully immature population or the heterogeneous age population). This reduction was implemented by scaling the afferent drive in a manner that was reliant on the neuronal diameter, with lower diameter translating to larger reduction in the synaptic drive, and was adjusted towards the goal of reducing firing rate variability across the neuronal population (Mishra and Narayanan, 2019).

### Network Analyses: Virtual Animal Traversal and Assessment of Channel Decorrelation in Networks Lacking Specific Ion Channels

A virtual animal was allowed to traverse a 1 m × 1 m arena, and the *x* and *y* coordinates of the animal’s location translated to changes in the external inputs from the MEC and LEC. The direction (range: 0–360°) and distance per time step (velocity: 2.5–3.5 m/s) were randomly generated, and were updated every millisecond. The amount of time taken for the virtual animal to approximately cover the entire arena was ∼1000 seconds. All simulations were performed for 1000 s, with the spatiotemporal sequence of the traversal maintained across simulations to allow direct comparisons, with the initial position set at the center of the arena. After the network was constructed with different forms of heterogeneities and/or with different ion channels knocked out, the spike timings of each GC were recorded through the total traversal period of 1000 s (Mishra and Narayanan, 2019). Note that the impact of knocking out the spike generating conductances (from either BC or GC) was not assessed because firing rate or response correlation could not be computed without a spiking response from the neurons.

The overall firing rate of granule cells (*e.g.*, Fig. 6*A*) spanning the 1000 s period was computed as the ratio between the spike count during the period and the total time (1000 s). For each network configuration with any of the 9 ion channel (7 from GCs and 2 from BCs) virtual knockouts, we computed change in this overall GC firing rate after virtual knockout of the channel. Quantitatively, let *F*_n_ (*base*) represent the overall firing rate of neuron *n* (1 ≤ *n* ≤ 100 for GC; 1 ≤ *n* ≤ 15 for BC) in the base version of the network, and let *F*_n_ (*C*_i_) represent the firing rate of the same neuron, obtained from a network where one of the channels *C*_i_ (1 ≤ *i* ≤ 7 for GC VKMs; 1 ≤ *i* ≤ 2 for BC VKMs) was knocked out from all neurons (in either the GC or BC population). We quantified the impact of single ion channel knockout on firing as a difference:

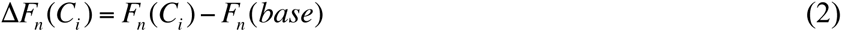

This procedure was repeated for different network configurations endowed with different sets of heterogeneities. The statistics of these measurements were plotted as quartiles to depict the entire span of changes (*e.g.*, Fig. 6*A*).

Instantaneous firing rates for each GC was computed from binarized spike time sequences by convolving them with a Gaussian kernel with a default standard deviation (σ_FR_) of 50 ms. Although all correlation computations (*e.g.*, Fig. 6*B*) were computed using a σ_FR_ of 50 ms, for displaying firing rate overlaid on the arena, we employed a smoother instantaneous firing rate computed with a σ_FR_ of 2 s (*e.g.*, Fig. 4*A*). This is important, because we had demonstrated that the choice σ_FR_ plays a critical role in correlation computation (Mishra and Narayanan, 2019).

**Figure 4.**
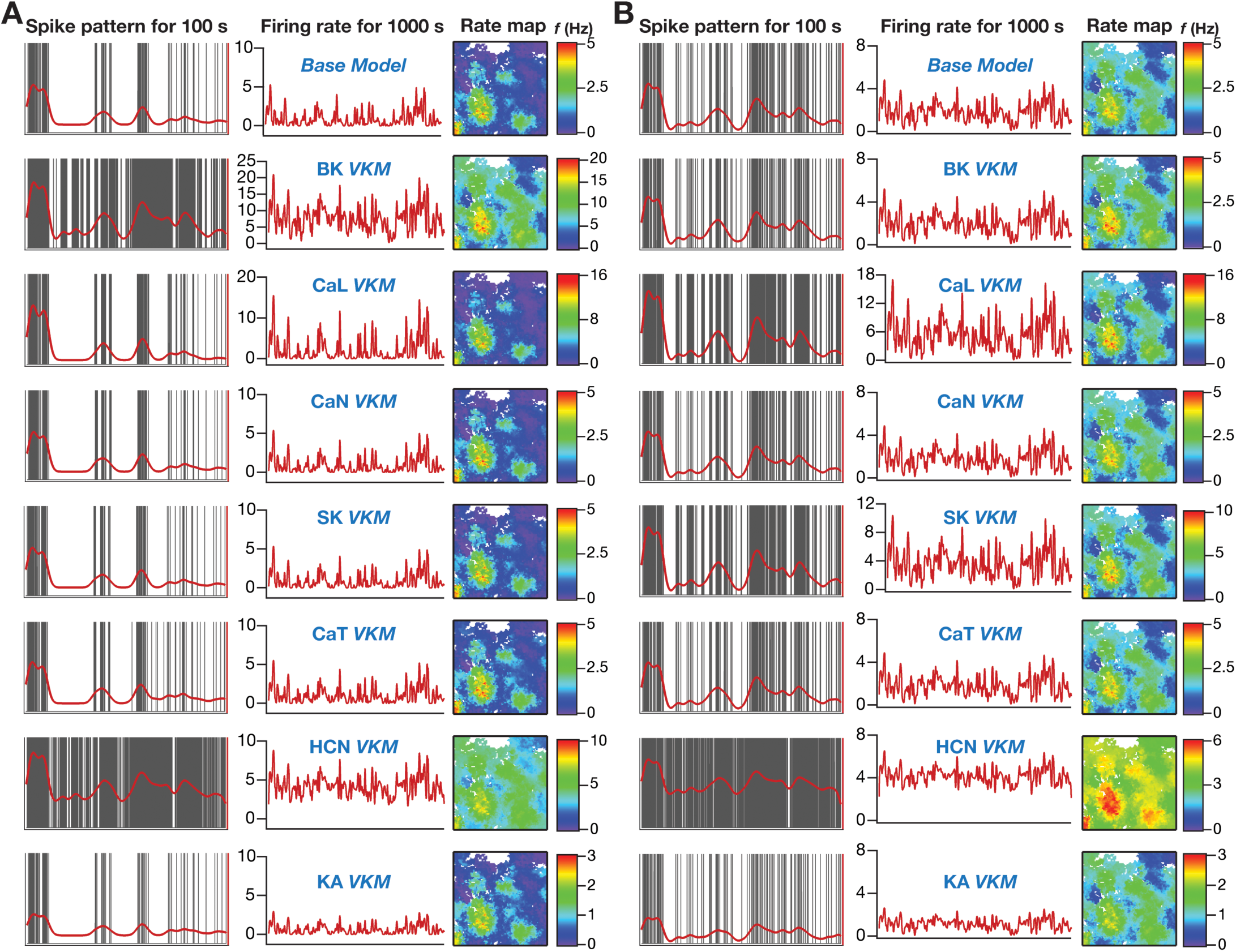
Granule cell firing profiles and spatial maps depicting the heterogenous impact of virtually knocking out individual ion channels from granule cells in a network receiving *identical* afferent inputs. *A*, *Left:* Spike patterns (gray) overlaid with firing rates (red) for a 100 s period for valid GC model 50, residing in a GC-BC network endowed with intrinsic and synaptic heterogeneities and receiving identical afferent inputs. *Center:* Instantaneous firing rates of GC model 50 for the entire 1000 s of animal traversal across the arena. *Right:* Color-coded spatial rate maps showing firing rate of GC model 50 superimposed on the trajectory of the virtual animal. The top-most panels represents these measurements for the base model (where all ion channels are intact), and the other panels depict these measurements obtained after virtual knockout of individual ion channels from the granule cell population of the network. *B,* Same as (*A*) for GC model 44 residing in the same network. Models 50 and 44 respectively showed maximum and minimum changes in firing rate after virtual knockout of BK ion channel (see Fig. 6A).

**Figure 5.**
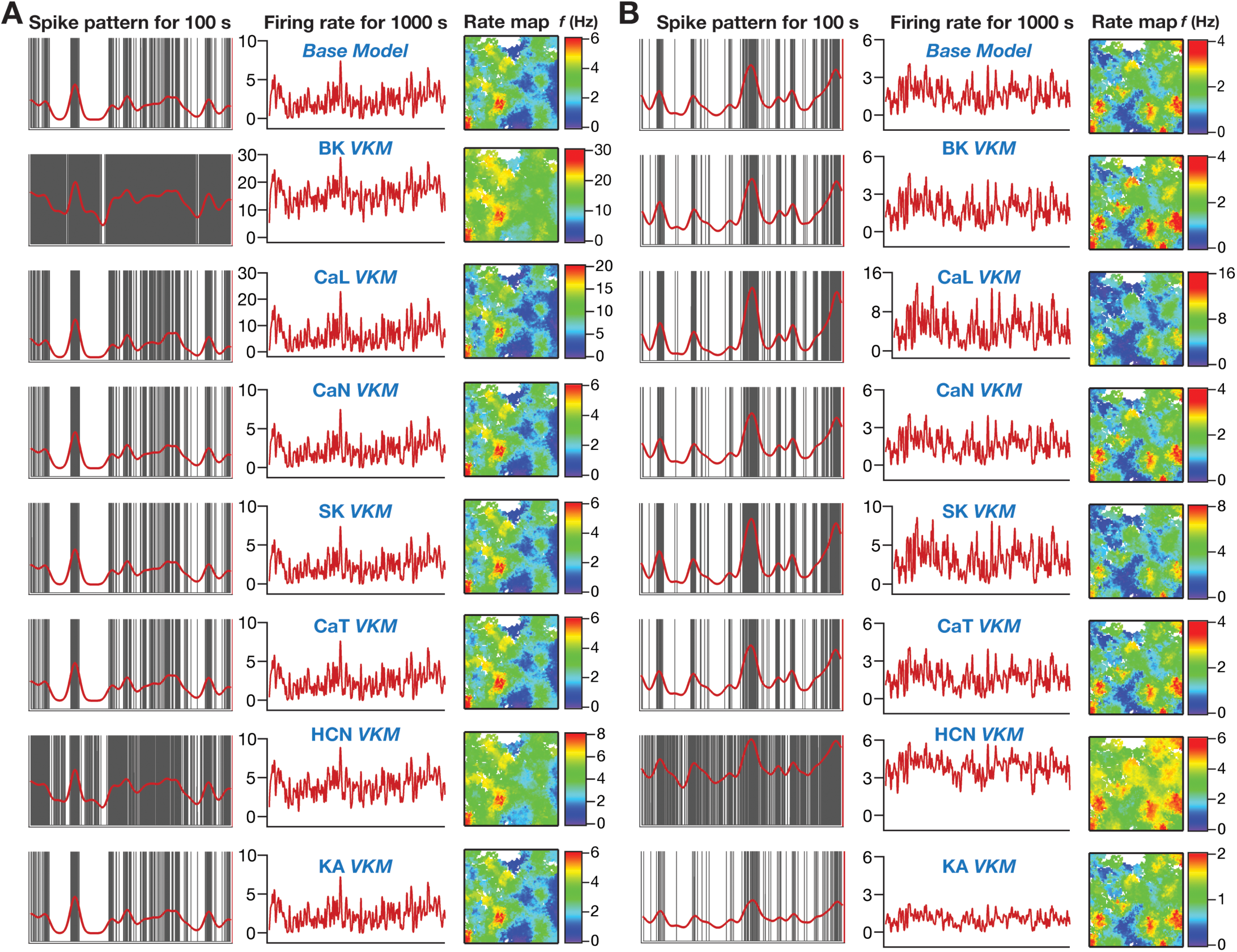
Granule cell firing profiles and spatial maps depicting the heterogenous impact of virtually knocking out individual ion channels from granule cells in a network receiving *heterogeneous* afferent inputs. *A*, *Left:* Spike patterns (gray) overlaid with firing rates (red) for a 100 s period for valid GC model 84, residing in a GC-BC network endowed with intrinsic and synaptic heterogeneities and receiving identical afferent inputs. *Center:* Instantaneous firing rates of GC model 84 for the entire 1000 s of animal traversal across the arena. *Right:* Color-coded spatial rate maps showing firing rate of GC model 84 superimposed on the trajectory of the virtual animal. The top-most panels represents these measurements for the base model (where all ion channels are intact), and the other panels depict these measurements obtained after virtual knockout of individual ion channels from the granule cell population of the network. *B,* Same as (*A*) for GC model 44 residing in the same network. Models 84 and 44 respectively showed maximum and minimum changes in firing rate after virtual knockout of BK ion channel (see Fig. 7A).

**Figure 6.**
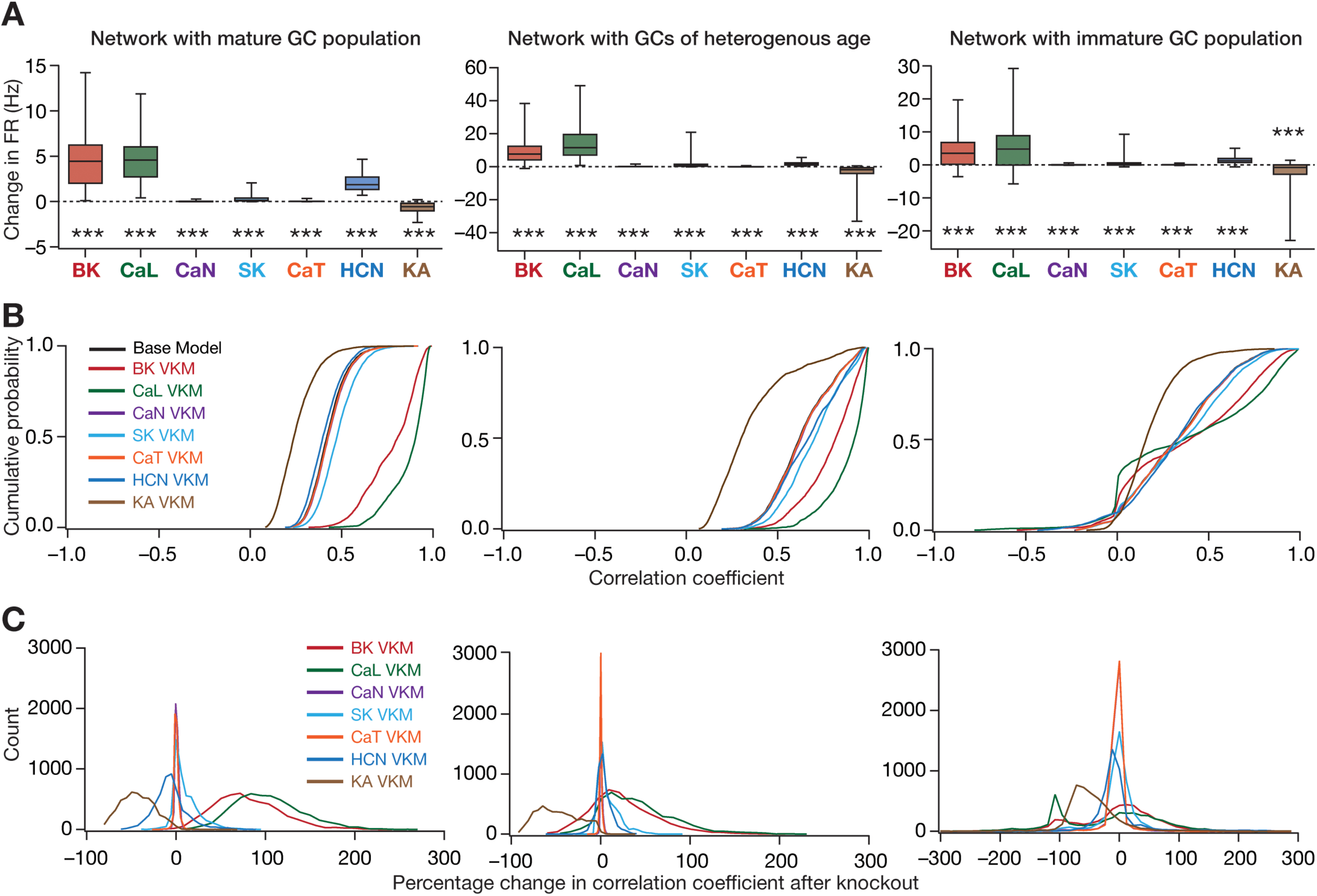
Virtual knockout of individual ion channels from granule cells resulted in differential and variable impact on channel decorrelation in networks endowed with distinct heterogeneities and receiving *identical* afferent inputs. *A*, Difference in firing rates (Eq. 2) for all granule cells in the network, represented as quartiles. Firing rate of each cell was computed from the spike count of the cell for the entire 1000 s traversal of the virtual animal. *p* values were obtained using Wilcoxon signed rank test, where the change in firing rate was tested for significance from a “no change” scenario. ***: *p* < 0.001. *B,* Cumulative distribution of inter-neuronal pairwise firing rate correlation coefficients for networks built with either the base models, or with models after virtual knockout of individual ion channels. Shown are plots for the base model network and for the networks built with GC neurons where one of the 7 ion channels was virtually knocked out. *C,* Distribution of percentage changes in correlation coefficients for neuronal responses from the VKM network, compared to the respective base model coefficients. Shown are plots corresponding to percentage changes in networks built with GC neurons where one of the 7 ion channels was virtually knocked out. For (*A–C*), plots are shown for simulations performed with three distinct networks and associated virtual knockouts: network with a fully mature GC population (*left*), network with a GC population of heterogeneous age (*center*) and network with a fully immature GC population (*right*). Note that all three networks are endowed with intrinsic and synaptic heterogeneities. All neurons in the network received identical afferent inputs.

Histograms of Pearson’s pairwise correlation coefficients were computed between instantaneous firing rate arrays (each spanning the 1000 s period) of each GC. Specifically, a correlation coefficient matrix was constructed, with the (*i*, *j*)^th^ element of this matrix assigned to the Pearson’s correlation coefficient (*R*_ij_) computed between the instantaneous firing rate arrays of neuron *i* and neuron *j* in the network (channel decorrelation). As these correlation matrices are symmetric with all diagonal elements set to unity, we employed elements in the lower triangular part for computing the associated histogram (*e.g.*, Fig. 7*B*). Note that in this study, our focus is on channel decorrelation, a form of response decorrelation that is distinct from pattern decorrelation, where the focus in on correlation between temporally-aligned response profiles of individual channels of information (*i.e.,* neurons) to afferent stimuli. Channel decorrelation is postulated to decrease the overlap between channel responses, resulting in a code that is efficient because the information conveyed by different channels is largely complementary. This is distinct from pattern decorrelation, which is assessed by computing response correlations across these two sets of neuronal outputs when inputs corresponding to two different patterns arrive onto the same network. Pattern decorrelation is computed to determine the ability of neuronal outputs to distinguish (pattern separation) between the two input patterns (Wiechert et al., 2010; Mishra and Narayanan, 2019).

**Figure 7.**
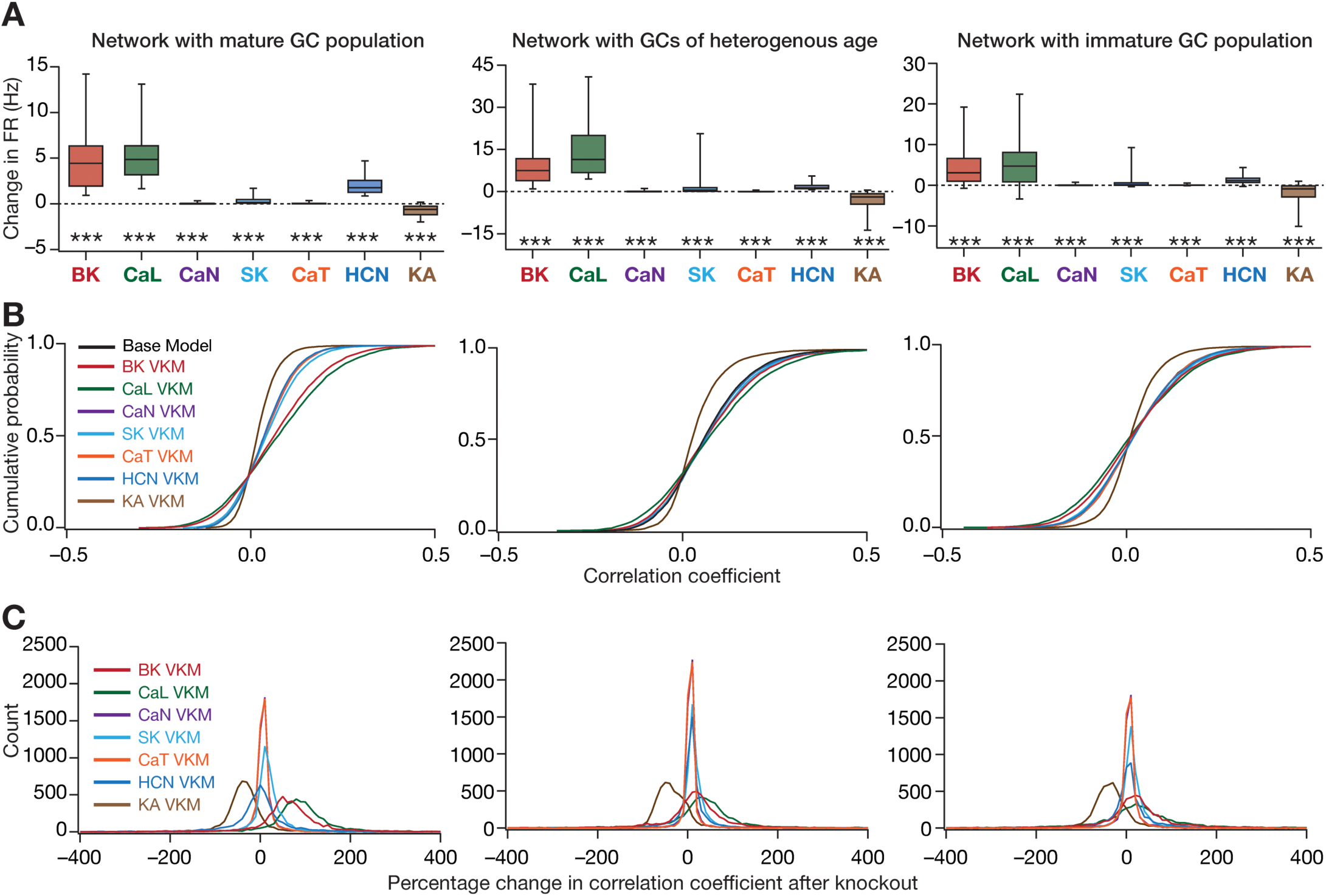
Virtual knockout of individual ion channels from granule cells resulted in differential and variable impact on channel decorrelation in networks endowed with distinct heterogeneities and receiving *heterogeneous* afferent inputs. *A*, Difference in firing rates (Eq. 2) for all granule cells in the network, represented as quartiles. Firing rate of each cell was computed from the spike count of the cell for the entire 1000 s traversal of the virtual animal. *p* values were obtained using Wilcoxon signed rank test, where the change in firing rate was tested for significance from a “no change” scenario. ***: *p* < 0.001. *B,* Cumulative distribution of inter-neuronal pairwise firing rate correlation coefficients for networks built with either the base models, or with models after virtual knockout of individual ion channels. Shown are plots for the base model network and for the networks built with GC neurons where one of the 7 ion channels was virtually knocked out. *C,* Distribution of percentage changes in correlation coefficients for neuronal responses from the VKM network, compared to the respective base model coefficients. Shown are plots corresponding to percentage changes in networks built with GC neurons where one of the 7 ion channels was virtually knocked out. For (*A–C*), plots are shown for simulations performed with three distinct networks and associated virtual knockouts: network with a fully mature GC population (*left*), network with a GC population of heterogeneous age (*center*) and network with a fully immature GC population (*right*). Note that all three networks are endowed with intrinsic and synaptic heterogeneities. Neurons in the network received heterogeneous afferent inputs.

In assessing channel decorrelation as a function of input correlation (Mishra and Narayanan, 2019), we first computed the total afferent current impinging on each neuron. As the total current was the same for scenarios where identical afferent inputs were presented, the input correlation across all neurons was set at unity. For the scenario where the afferent inputs were heterogeneous, pairwise Pearson correlation coefficients were computed for currents impinging on different DG neurons and were plotted against the corresponding response correlation (for the same pair). Output correlations in this plot were binned for different values of input correlation, and the statistics (mean ± SEM) of response correlation were plotted against their respective input correlation bins (*e.g.*, Fig. 8*A*).

**Figure 8.**
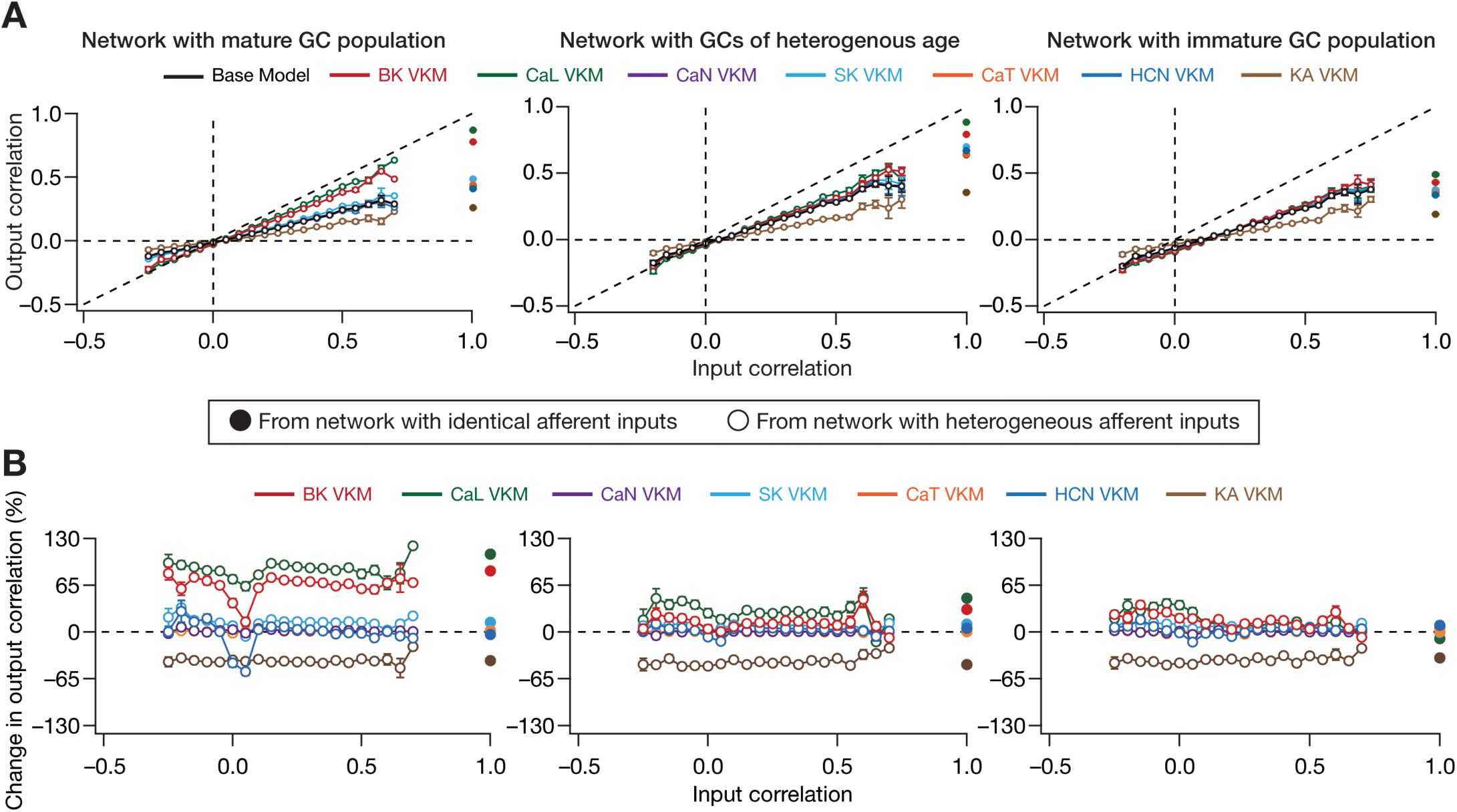
The impact of virtual knockout of individual ion channels on the degree of channel decorrelation achieved by a network depends on the specific ion channel being knocked out, and is independent of age and afferent heterogeneities. *A*, Pairwise response (output) correlation plotted as a function of the corresponding pairwise input correlation, for the base model network and for networks built with GC neurons where one of the 7 ion channels was virtually knocked out. *B,* Percentage change in response (output) decorrelation in VKM networks with reference to the corresponding response decorrelation in the base model, plotted as functions of input correlation. For (*A–B*), plots are shown for simulations performed with three distinct networks and associated virtual knockouts: network with a fully mature GC population (*left*), network with a GC population of heterogeneous age (*center*) and network with a fully immature GC population (*right*). Note that all three networks are endowed with intrinsic and synaptic heterogeneities. In all cases, network outcomes are represented as solid or open circles, when the network received identical or heterogeneous afferent input, respectively. Note that the input correlation is unity for networks receiving identical inputs, whereas input correlation is dependent on specific pairs of inputs when the network receives heterogeneous inputs.

To compute pairwise changes in the degree of decorrelation consequent to channel knockout, we calculated percentage changes in the specific pairwise Pearson’s correlation coefficient after knockout of a specific channel, compared to the coefficient’s value before knockout. Quantitatively, let *R*_ij_ (*base*) represent the Pearson’s correlation coefficient computed between the instantaneous firing rate arrays of neuron *i* and neuron *j* (1 ≤ *i, j* ≤ 100; *i* ≠ *j*) in the base version of the network. Let *R*_ij_ (*C*_k_) represent the Pearson’s correlation coefficient computed between the same neuronal pair (*i*, *j*), obtained from a network where one of the channels *C*_k_ (1 ≤ *k* ≤ 7 for GC VKMs) was knocked out from all neurons (in either the GC or BC population). We quantified the impact of single ion channel knockout on output correlation of GCs as a percentage change:

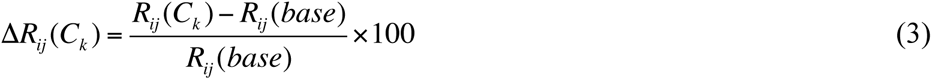

This procedure was repeated for different network configurations endowed with different sets of heterogeneities. The statistics of these percentage changes were plotted as histograms (*e.g.*, Fig. 6*C*), or were plotted against their respective input correlation values that were binned (*e.g.*, Fig. 8*B*) using a procedure similar to the output correlation *vs.* input correlation plot mentioned above.

### Computational details

All simulations were performed using the NEURON simulation environment (Carnevale and Hines, 2006), at 34° C with an integration time step of 25 µs. Analysis was performed using custom-built software written in Igor Pro programming environment (Wavemetrics). Statistical tests were performed in statistical computing language R (www.R-project.org).

## RESULTS

The primary objective of this study was to address the question on whether and how individual ion channels alter channel decorrelation in a DG network endowed with distinct forms of biological heterogeneities? The very nature of the question involved traversal across three distinct scales of analyses (ion channels-neurons-network), and required that we account for the different biological heterogeneities expressed in the DG. Therefore, we built a multi-scale DG network, which was constructed from a heterogeneous population of biophysically constrained and electrophysiologically validated conductance-based neuronal models for both GCs (Fig. 1*A*) and BCs (Fig. 1*B*). The afferent inputs to the DG network from the medial and lateral entorhinal cortices were driven by the position of a virtual animal traversing a 1 m × 1 m arena (Fig. 1*C*). We constrained the local excitatory-inhibitory connectivity and scaled the afferent inputs from the EC such that the firing rates of GCs and BCs matched their electrophysiological counterparts from *in vivo* recordings from awake-behaving animals. We recorded firing rates from all granule cells within the network and employed them for further analyses. This configuration provided us with an ideal setup to understand the impact of components in the molecular scale (ion channels) on functional outcomes (response decorrelation) at the network scale, after rigorously accounting for cellular-scale physiological properties and how they emerge from interactions among disparate channels expressed at the molecular scale. We also employed appropriate techniques developed earlier (Mishra and Narayanan, 2019) to incorporate four distinct forms of heterogeneities into this network (Fig. 1*D*), to analyze the impact of individual ion channels on response decorrelation in networks configured with different sets of these heterogeneities. As a first step in our analyses, we assessed the impact of virtually knocking out individual ion channels on single-neuron properties of the heterogeneous GC and BC populations.

### Multiple ion channels differentially impact different single-neuron physiological features of granule and basket cells

The variable expression of a plethora of ion channels in individual neuronal subtypes, along with the interactional complexity that spans the different subunits of these ion channels, the signaling molecules that alter their expression and function, and structural/functional interactions across different ion channels, bestows a neuron with its signature physiological characteristics. Whereas certain ion channels play a dominant role in *mediating* specific sub- and supra-threshold properties of neuron (*e.g.,* action potential generation through an interplay between fast sodium and delayed-rectifier potassium ion channels, spike frequency adaptation emerging through calcium-mediated interactions between calcium and calcium-dependent potassium ion channels), others play a regulatory role by *modulating* or *refining* fine details of neuronal physiology (*e.g*., modulation of neuronal firing rate and afterhyperpolarization potentials by transient potassium ion channels, modulation of action potential width by repolarizing ion channels). The heterogeneous populations of GCs and BCs were arrived through independent and unbiased stochastic searches, employing ion channels characterized from these neurons and physiologically constrained by nine different physiological measurements, again measured from these neuronal subtypes. How do the 9 different active ion channels expressed in GCs and the 4 different active ion channels in BCs impact their respective physiological measurements?

To understand the contribution of these VGICs on single-neuron physiology of the heterogeneous populations GCs and BCs, each independently exhibiting ion-channel degeneracy (Mishra and Narayanan, 2019), we turned to virtual knockout models (Rathour and Narayanan, 2014; Anirudhan and Narayanan, 2015; Mukunda and Narayanan, 2017; Basak and Narayanan, 2018). Here, each conductance in a given model was independently set to zero, and the 9 single-cell physiological measurements were recomputed after this virtual knockout of the ion channel. Examples of virtual knocking out each of the 9 ion channels in 4 different GC models are depicted in Figure 2. These examples illustrate the differential and variable response of individual models to ion channel knockouts. Firstly, considering the example model #36 (Fig. 2*A*), we noticed that the changes observed in the sub-threshold (input resistance) and supra-threshold (firing rate for 150 pA pulse current injection) responses were *differential*, with reference to knocking out different ion channels. For instance, knocking out the CaN ion channels did not change the input resistance or firing rate, knocking out BK ion channels introduced large changes to input resistance and firing rate, but knocking out either NaF or KDR ion channels altered firing rate without altering input resistance. These observations pointed to different measurements in the *same* model being *differentially* sensitive to different ion channel knockouts. Second, valid model #36 (Fig. 2*A*) and #50 (Fig. 2*B*), respectively represent the minimum (100×(231–214)/214=7.9%) and maximum (100×(287–156)/156=83.9%) percentage change in *R*_in_ after virtual knockout of HCN ion channels. Thus, whereas the contribution of HCN ion channels to *R*_in_ is low for model #36, the contribution is higher for model #50. However, the contribution of BK ion channels to *R*_in_ is high for model #36, whereas it is lower for model #50. This represents the *variability* in the dependence of different models on the *same* ion channel in regulating a given physiological measurement, and demonstrates that the contribution of a given structural component to a functional measurement is heterogeneous (Rathour and Narayanan, 2019).

This observation is further emphasized by valid model #100 (Fig. 2*C*) and #41 (Fig. 2*D*), respectively representing the minimum (100×(22–15)/15=46.6%) and maximum (100×(60–11)/11=445%) percentage change in *f*_150_ after virtual knockout of *L-*type calcium ion channels. The differential and variable response of neuronal models to ion channel knockouts might be observed from the four examples shown here. These examples emphasize the need to assess models employing unbiased stochastic searches, and the need to account for the heterogeneities inherent to neuronal populations. If a hand-tuned model, which let’s say arrives at parameters that are close to one of these four models, the conclusions would be based *solely* on the ion channel composition in that single hand-tuned model, arriving at biased conclusions about the role of specific ion channels in regulating specific measurements across the entire population of neurons. The heterogeneous population arrived employing the unbiased stochastic search and the VKM approach together enabled recognition and quantification of the differential and variable dependence of different measurements on distinct ion channels in disparate models.

We repeated VKMs for all the 126 models and 9 ion channels in GC population (Fig. 1*A*), and the 54 models and the 4 ion channels in BC population (Fig. 1*B*), spanning all the 9 physiological measurements for both populations (Fig. 3). As NaF and KDR ion channels didn’t alter sub-threshold measurements significantly, and the absence of either of these ion channels resulted in loss of action potential firing or repolarization, we have not included these VKM results. In addition, as the firing rate of the neuron in response to 50 pA current changed only for a small proportion of models and knockouts, we have not incorporated that measurement into our results. Therefore, we analyzed 8 measurements each from VKMs for 7 ion channels in the GC population, and VKMs for 2 ion channels in the BC population (Fig. 3). In what follows, we systematically analyze the impact of each ion channel subclass on the different physiological measurements, from a population perspective for both GCs and BCs.

Our granule cell model expressed three voltage-gated calcium ion channels. We noted that the elimination of the non-inactivating CaL ion channels had the largest impact on ISI ratio (Fig. 3*E*) and firing rate (Fig. 3*F*), but showed very little effect on both subthreshold measurements (Figs. 3*G–H*). There was significant variability on how these ion channels altered firing rate and ISI ratio on different models (Figs. 3*E–F*). Elimination of the inactivating *N*-type calcium ion channels did not introduce large changes in any of the sub- or supra-threshold measurements considered here, and the variability across models was lower as compared to the variability across CaL VKMs. VKMs of the low-voltage activated inactivating CaT ion channels exhibited small and variable changes in AP threshold (Fig. 3*C*), fast afterhyperpolarization (Fig. 3*D*), ISI ratio (Fig. 3*E*) and firing rate (Fig. 3*F*). In addition, given their low-voltage activation, CaT ion channels also impacted input resistance to a small extent (Fig. 3*G*).

Of the two voltage-gated potassium ion channels incorporated into GC models, the absence of the non-inactivating KDR ion channels resulted in improper repolarization of action potentials, and did not considerably alter sub-threshold properties (*e.g.*, Fig. 2). Lack of the transient KA VKMs significantly altered all supra-threshold measurements (Figs. 3*A–F*), with greater impact on AP amplitude (Fig. 3*A*) and AP half-width (Fig. 3*B*), also manifesting significant variability across models (Figs. 3*A–F*). Importantly, although elimination of a potassium ion channel is generally expected to *increase* excitability, in a large proportion of models, we observed a counterintuitive *reduction* in firing rate after virtual knockout of KA ion channels (Fig. 3*F*). However, such counterintuitive changes to firing rates have been explained through *functional interactions* across ion channels (Kispersky et al., 2012), and the interactions between KA and other repolarizing channels have been shown to explain similar counter-intuitive results observed with changes in KA ion channel conductances (Narayanan and Johnston, 2010; Anirudhan and Narayanan, 2015). In our case, the explanation emerges from the impact of KA ion channels on other measurements (Fig. 3). Specifically, in KA VKMs, the AP amplitude (Fig. 3*A*) and AP half width (Fig. 3*B*) are considerably larger, implying a higher degree of voltage-dependent activation of KDR channels and a larger influx of calcium through the VGCCs, resulting from the large-amplitude and wide action potentials. This, in turns, results in a larger fraction of these other repolarizing ion channels (KDR, SK and BK) opening, reflected in a larger afterhyperpolarization (Fig. 3*D*) and a higher adaptation (Fig. 3*E*). The large afterhyperpolarization leads to a longer time for the membrane to charge up to threshold, and together with the higher adaptation resulted in the observed reduction in firing rates in the large proportion of models. Thus, the relative dominance of other ion channels in the repolarization kinetics and the enhanced action potential amplitude in the absence of KA ion channels explains the counterintuitive reduction in firing rate in their VKMs. As expected from the inactivating nature of these ion channels and the hyperpolarized resting potentials of GCs, KA ion channels did not significantly affect subthreshold measurements (Fig. 3*G–H*).

Turning to calcium-activated potassium ion channels, the VKMs of either BK or SK ion channels showed the largest and highly variable impact on ISI ratio (Fig. 3*E*) and on firing rate (Fig. 3*F*), having relatively weaker impact on the other sub and supra threshold measurements. Based on the similarity in the outcomes of knocking out CaL or the calcium-activated potassium channels on physiological responses, we reasoned the increase in excitability after CaL knockouts to be consequent to their interactions with the calcium-dependent potassium ion channels. The lack of hyperpolarization-activated cyclic nucleotide gated ion channels (HCN or *h*) introduced large and variable changes to input resistance (Fig. 3*G*) and sag ratio (Fig. 3*H*), given its expression at rest as well as its hyperpolarization-induced activation profile. This robust impact on sub-threshold properties of neuron also translated to mild changes to other supra threshold measurements (Figs. 3*A–F*).

In the BC population, very similar to their counterparts in the GC population, HCN ion channels dominantly influenced sub-threshold measurements (Fig. 3*G–H*). However, in contrast to majority of the GC models, here VKMs of KA ion channels exhibited strong *increase* in firing rate (Fig. 3*F*), which was consistent with a lack of change in AP amplitude (Fig. 3*A*) and half width (Fig. 3*B*) in KA VKMs of these neurons. Together, our analyses of the impact of different ion channels on single-neuron physiology of the heterogeneous GC and BC populations demonstrated differential and variable dependence of the various physiological measurements on these ion channels. Importantly, the mapping between ion channels and physiological measurements was many-to-many, where different ion channels affected any given measurement, and any specific ion channel altered several measurements.

### Virtual knockout of individual ion channels across GCs introduced differential and variable scaling of firing rate profiles and associated spatial maps

Our analysis thus far was focused on the expression and perpetual interactions amongst multiple ion channels that underlie single neuronal functions. Akin to such cellular-scale interactions, microcircuit scale interactions occur to produce specific network outputs, synergistically integrating the intrinsic attributes of individual neurons along with synaptic information that arrives from other neurons within or external to the local network. Motivated by the differential and variable impact of individual ion channel virtual knockouts on single neuronal measurements, we next asked how these individual channel knockouts from GCs affect their firing rate profiles and their spatial maps with reference to the virtual animal traversal (Fig. 1*C*). In performing VKM simulations, everything else in the network was set identical to baseline conditions, including the specific spatio-temporal trajectory of the virtual animal in the arena, except for knocking out one specific ion channel from all GCs in the network. The procedure was repeated for the 7 GC ion channels, and the firing rates of networks built with the VKMs were compared with those of the network with base models.

The impact of virtually knocking out each of the 7 GC ion channels on firing rate profiles and spatial map profiles of four different example GCs residing in two distinct networks are shown in Figures 4–5. Whereas Figure 4 shows these profiles for a network receiving *identical* afferent inputs from the EC, Figure 5 depicts profiles for a network endowed with *afferent heterogeneities*, whereby each neuron in the network received distinct sets of EC inputs. The networks employed in these illustrative examples (Fig. 4–5) were endowed with intrinsic as well as synaptic heterogeneities, but did not express neurogenesis-induced structural heterogeneities.

Reminiscent of our observations with single-neuron physiological measurements, we found that differential impact of different ion channel knockouts on the profiles of the same neuron, and variable impact of knocking out the same ion channel on different neurons within the same network. For instance, in Figure 4, in GC #50 (Fig. 4*A*), deletion of either HCN or BK ion channels resulted in large increases in firing rate, whereas KA VKMs exhibited reductions in firing rates and the other knockouts did not elicit significant differences in firing rate profiles across space. With reference to variability in the impact of knockouts, in GC #44 within the same network (Fig. 4*B*) recorded during the same virtual traversal, BK, CaL or CaT VKMs did not have a significant effect, but deletion of CaL, SK or HCN ion channels resulted in increased firing rates and KA VKMs again exhibited reduced firing rates. The firing rate changes were introduced by knockouts belonged to two broad categories: multiplicative scaling, where changes were restricted to the locations where the base model fired (*e.g.*, SK VKM of GC #44 compared to its baseline of Fig. 4*B*), and additive scaling, where there was a shift in the entire firing profile (*e.g.*, HCN VKM of GC #44 compared to its baseline of Fig. 4*B*). In some cases, we observed a combination of multiplicative and additive scaling (*e.g.*, BK VKM of GC #50 compared to its baseline of Fig. 4*A*). As the network depicted in Fig. 4 received *identical* EC inputs, it may be noted that the place field locations were identical across the two cells, with differences only in firing rates between these two cells and across knockouts.

We confirmed these observations to also extend to a network endowed with afferent heterogeneities (Fig. 5). Specifically, the differential (*e.g.*, SK *vs.* BK VKMs of GC #84 in Fig. 5*A*) and variable (*e.g.*, BK VKMs of GC #84 in Fig. 5*A* *vs.* GC #44 in Fig. 5*B*) responses to ion channel knockouts in the same set of neurons within the same network during the same virtual traversal were observed. Firing rate changes were scaled either multiplicatively (*e.g.*, KA VKMs of GC #44 in Fig. 5*B*) or additively (*e.g.*, HCN VKMs of GC #44 in Fig. 5*B*) or both (*e.g.*, BK VKMs of GC #44 in Fig. 5*A*). However, as the network depicted in Fig. 5 received heterogeneous EC inputs, the place field locations were different between the two cells within the same network during the same virtual traversal. Virtual knockouts, however, only scaled the firing rates of individual cells, without altering the position of place fields where a given cell was firing (*e.g.*, compare the firing profiles and spatial maps of GC #84 in Fig. 5*A* under baseline conditions and in the VKM network).

We quantified these variable and differential responses of neural firing to the virtual knockout of different ion channels across the entire population of GCs for network receiving *identical* (Fig. 6*A*) or heterogeneous (Fig. 7*A*) EC inputs, with different degrees of structural heterogeneity. With reference to degrees of structural heterogeneities, we employed three configurations (Mishra and Narayanan, 2019): two networks with fully mature or fully immature GC populations, which did not have any structural heterogeneities and a network that was endowed with a heterogeneous structural properties (Fig. 6, Fig. 7). Whereas the fully mature population refers to a scenario where there are no new neurons integrated into the circuit, the fully immature population is an artificial setting where all neurons are immature with less surface area and the heterogeneous population is reflective of a more natural milieu where neurons are at different stage of maturation. Although the quantitative changes in firing rate were dependent also on the specific kind of structural heterogeneities expressed in the network, the direction and strength of changes in GC firing rate in a network were similar to those at the single-neuron scale (compare Fig. 6*A* and Fig. 7*A* with Fig. 3*F*). Specifically, in a large proportion of GCs, knockout of CaL or BK ion channels introduced large increases in firing rate, elimination of CaN or CaT or HCN or SK ion channels resulted in relatively smaller increases in firing rate, and KA ion channel VKMs exhibited relatively small reduction in firing rate compared to their respective base models. Interpretations of changes, however, should not be drawn from the summary statistics, but should be driven by heterogeneities (Rathour and Narayanan, 2019). It should be noted that there are significant differences in different neurons in the same network during the same virtual traversal in terms of which ion channel plays a dominant role in altering firing rate.

Quantitatively, although the strength of afferent drive was scaled depending on the maturity (surface area) of the neuron to ensure that network firing rates were comparable across the three networks, VKMs resulted in larger changes in firing rates of neurons endowed with immature neurons. This was irrespective of whether the network received *identical* (Fig. 6*A*) or heterogeneous (Fig. 7*A*) afferent inputs. This should be expected because of the higher excitability of the relatively immature neurons, whereby even smaller changes to currents (here due to loss of specific ion channels) result in larger changes to voltage responses. Together, virtual knockout of individual ion channels across GCs resulted in differential and variable scaling of firing rate profiles and associated spatial maps, with larger changes observed in networks where immature neurons were present.

### The impact of eliminating individual ion channels on channel decorrelation was differential and variable in networks endowed with different heterogeneities

The location of the DG, its unique features of sparse and diverse connectivity in conjunction with the expression of adult neurogenesis has led to postulates of its role in response decorrelation and pattern separation. Channel decorrelation, one form of decorrelation of network responses, is assessed by computing pair-wise correlations across temporally aligned outputs of individual neurons (channels) within the network, when inputs corresponding to a single virtual arena traversal arrive onto the network. We had earlier established that distinct forms of network heterogeneities could synergistically interact with each other in mediating channel decorrelation in the DG, apart from establishing a dominance hierarchy among these forms of heterogeneities when they co-expressed (Mishra and Narayanan, 2019). Does altering neuronal intrinsic properties through virtual knockout of individual ion channels regulate channel decorrelation in the DG network?

We plotted the distribution of pairwise correlation coefficients of firing rate profiles of the different GCs in the network, for the base network (where all channels were intact), and for each of the 7 GC VKMs. We repeated this procedure for scenarios where inputs from EC were *identical* (Fig. 6*B*) or heterogeneous (Fig. 7*B*), for different degrees of structural heterogeneity in each case. We found differential effects of knocking out different ion channels on channel decorrelation in these networks. Consistent with our previous observations (Mishra and Narayanan, 2019), we noted that the degree of decorrelation observed in networks with heterogeneous afferent inputs (Fig. 7*B*) was higher than networks with *identical* afferent inputs (Fig. 6*B*). We also confirmed prior observations (Mishra and Narayanan, 2019) that the impact of different structural heterogeneities on networks receiving *identical* EC inputs was higher than the impact on networks receiving heterogeneous EC inputs (compare the base model distributions of correlation coefficients across the three different forms of heterogeneities in Fig. 6*B* vs. Fig. 7*B*). However, qualitatively, the effects of knocking out different channels on channel decorrelation were consistent across different network configurations, each endowed with distinct afferent and structural heterogeneities (Fig. 6*B*, Fig. 7*B*).

Specifically, across the different network configurations, elimination of either BK or CaL ion channels from the GCs of the network resulted in large rightward shifts in the cumulative distribution of response correlation coefficients, indicative of a reduction in response decorrelation (Fig. 6*B*, Fig. 7*B*). On the other hand, virtual knockout of KA ion channels resulted in a significant leftward shift in the cumulative distribution of response correlation coefficients, indicating enhanced decorrelation in these networks. The other VKMs, of CaN, CaT, SK and HCN ion channels, resulted in relatively smaller shifts to the correlation coefficient distributions (Fig. 6*B*, Fig. 7*B*). These observations were further confirmed by the distributions of VKM-induced percentage changes in correlation coefficients for networks receiving *identical* (Fig. 6*C*) or heterogeneous (Fig. 7*C*) afferent inputs, and endowed with different degrees of structural heterogeneities.

### Networks endowed with high-excitability immature neurons manifested resilient channel decorrelation after virtual knockout of ion channels

How did the elimination of individual ion channels alter output response correlation as a function of different input correlations? Did ion channel knockouts specifically affect inputs with lower or higher correlation? Did the presence of neurogenesis-induced structural differences in the DG network alter the quantitative impact of different ion channel knockouts on channel decorrelation?

To address these questions, we placed output correlation coefficients of neuronal pairs into specific bins corresponding to their respective input correlation coefficients, and plotted the statistics of output correlations as functions of input correlations (Mishra and Narayanan, 2019). Whereas the network with *identical* afferent inputs had an input correlation of unity across all neuronal pairs, the network with heterogeneous afferent inputs was endowed with a range of pairwise input correlation coefficients depending on the specific nature of inputs that the neurons received (Fig. 8*A*). As established earlier (Mishra and Narayanan, 2019), the base network (where all channels were intact) manifested robust channel decorrelation, whereby the average output correlation coefficients were lesser than the average input correlation across all observed input correlations (Fig. 8*A*; black traces).

When we measured the output correlations as functions of their respective input correlations after elimination of individual GC channels, we made two important observations. First, in the presence of immature neurons, either in a network that was constructed entirely with immature neurons or in a network that was endowed with structural heterogeneities, we found that the amount of VKM-induced changes in channel decorrelation to be lesser compared to a network that was constructed of mature neurons (Fig. 8*A*). This was also reflected in the percentage changes observed in output correlation plotted as functions of input correlations (Fig. 8*B*). Second, for a given VKM, the average percentage change in output correlation was invariant to the average input correlation, across networks endowed with different structural and afferent heterogeneities (Fig. 8*B*). Although some VKMs elicited an enhancement and other introduced a reduction in the correlation coefficients, and although the quantitative changes were dependent on the specific heterogeneities expressed in the network (Fig. 6–8), the average percentage changes in output correlation was largely independent of the specific values of input correlation (Fig. 8*B*). This observation extended to output correlation coefficients measured for *identical* inputs (input correlation coefficient=1) as well (Fig. 8*B*).

### Virtual knockout of either the KA or the HCN ion channels from basket cells did not significantly alter GC firing rates or network decorrelation

Thus far, our analyses were confined to VKMs of GC neurons. What is the impact of altering ion channel composition in the basket cells of the network? We measured GC firing rates and decorrelation across GC responses in networks built with BCs lacking either the KA or the HCN ion channels (Fig. 9). We compared these outcomes with the base network (where all ion channels were intact), and found that elimination of either KA or HCN ion channels from BCs did not considerably alter GC firing rates across the network or introduce prominent changes in channel decorrelation computed across GC responses. This was consistent across networks with different configurations involving disparate combinations of structural and afferent heterogeneities (Fig. 9). We noted these to be simply a reflection of the relatively minor role played by the two ion channels in regulating BC intrinsic properties (Fig. 3).

**Figure 9.**
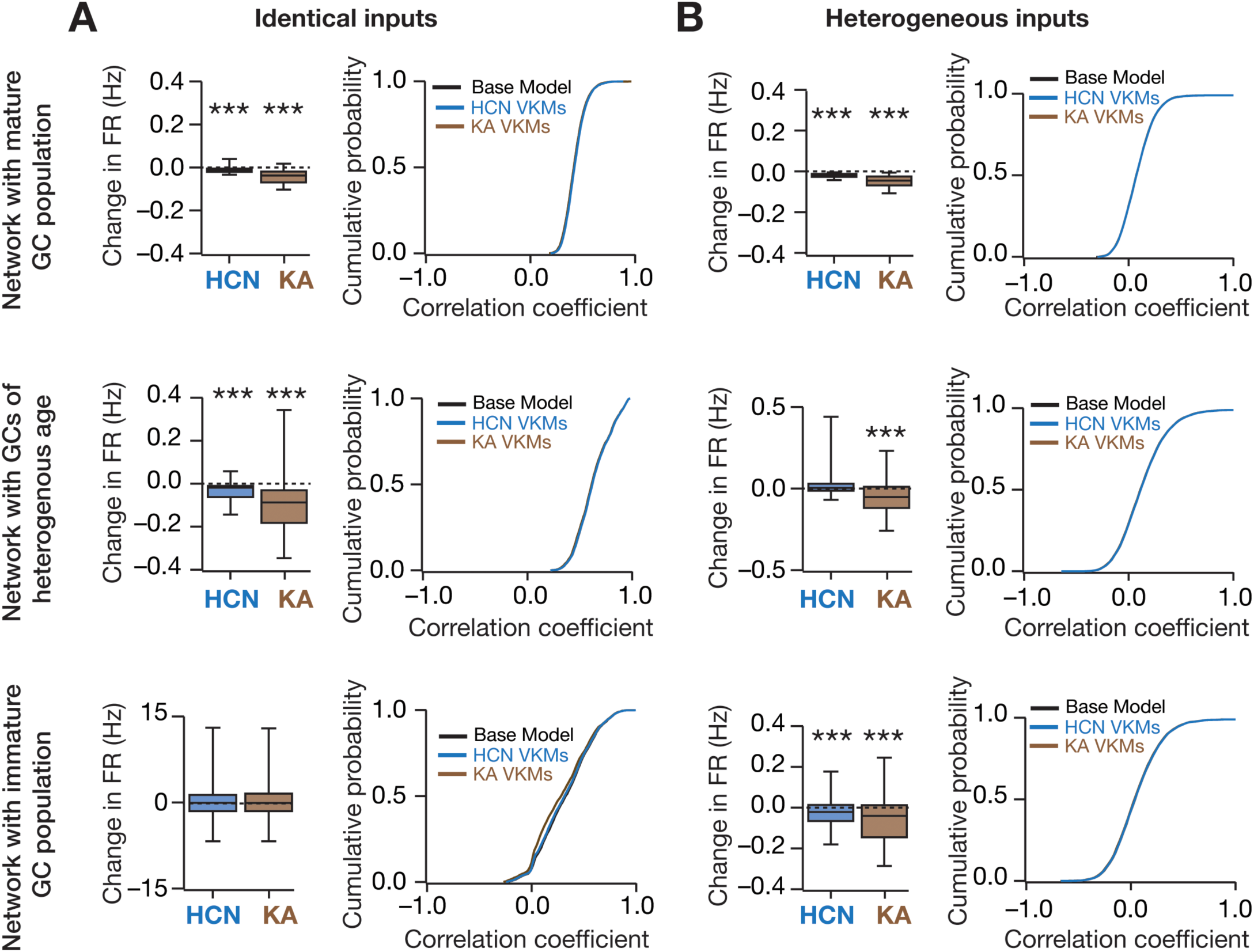
Weak impact of virtually knocking out HCN or KA ion channels from the basket cell population on firing rate and channel decorrelation in granule cell responses. *A*, *Top*: *Left,* Change in granule cell firing rate in a network receiving *identical* afferent input, represented as quartiles. Shown are the cases for the HCN or KA ion channel knockout from the basket cell population. *Right,* Cumulative distribution of granule cell firing rate correlation coefficient for different ion channel knockouts. Shown are the cases for the baseline network, and for cases where HCN or KA ion channels were virtually knocked out from the basket cell population. Plots in the other panels are similar to the top panel graphs, but for networks with different age heterogeneities: network with a fully mature GC population (*top*), network with a GC population of heterogeneous age (*middle*) and network with a fully immature GC population (*bottom*). Note that all three networks are endowed with intrinsic and synaptic heterogeneities. *B,* Same as (*A*) for network receiving heterogeneous afferent inputs. *p* values were obtained using Wilcoxon signed rank test, where the change in firing rate was tested for significance from a “no change” scenario. *** *p* < 0.001.

## DISCUSSION

In this study, employing systematic multi-scale analyses, we demonstrate differential and variable impacts of eliminating individual ion channels on single-neuron physiological properties, on network excitability and on channel decorrelation in DG networks configured with different forms of heterogeneities. At the single-neuron scale, our analyses, spanning both the GC and BC populations, revealed that the mapping between ion channels and physiological measurements was many-to-many, where different ion channels affected any given measurement, and any specific ion channel altered several measurements. At the network scale, our results demonstrated that individual ion channels expressed in granule cells play an important role in shaping DG network excitability, and critically regulate the ability of the network to perform channel decorrelation. The impact of knocking out individual ion channels was differential and variable both in terms of affecting network firing rates and channel decorrelation, but also was critically reliant on the specific local heterogeneities expressed in the DG network. Importantly, in the presence of structurally immature neurons in the DG network, the impact of ion channel elimination on channel decorrelation was considerably lower when compared with a network exclusively constructed with structurally mature neurons. These results highlight the role of local heterogeneities in regulating the resilience of the DG network to large network-wide perturbations to neuronal ion channel composition, even with the expression of the dominant afferent heterogeneities. Finally, we showed that for a given VKM, the average percentage change in output correlation was invariant to the specific values of input correlation, across different network configurations endowed with disparate structural and afferent heterogeneities.

### Heterogeneities in ion channel regulation of neuronal and network physiology

In our study, we report a many-to-many relationship between ion channels and neuronal and network physiology. What are the implications for this relationship, and the underlying variability in ion channel expression and in their ability to regulate neuronal and network physiology? The ubiquitously expressed abundance of variability in ion channel expression in each of the several neuronal subtypes imparts unique features to single neuron and network physiology. First, this variability forms the substrate for ion-channel degeneracy where similar physiological outcomes are achieved through disparate combinations of ion channels and their properties. This provides neurons with considerable flexibility in stitching together their signature physiological characteristics and in maintaining robustness in these properties, without strong constraints on the specific expression profiles of individual ion channels (Goldman et al., 2001; Marder and Goaillard, 2006; Grashow et al., 2010; Marder and Taylor, 2011; Kispersky et al., 2012; Rathour and Narayanan, 2012a, 2014; Drion et al., 2015; Basak and Narayanan, 2018; Mittal and Narayanan, 2018; Mishra and Narayanan, 2019; Rathour and Narayanan, 2019). Second, the variability in ion channel expression profiles implies that the response of neurons to even *identical* stimuli could be distinct, depending on the specific ion channels that are expressed across individual neurons, on the specific state of the neuron, on the nature of the afferent stimulus and how they activate/deactivate/inactivate the different ion channels, on the impact of different neuromodulators on these ion channels, and on specific activity-dependent plasticity profiles of these ion channels. This allows such intrinsic variability to form *a* substrate for decorrelating afferent stimuli (Padmanabhan and Urban, 2010; Mishra and Narayanan, 2019).

Third, depending on variable expression and the specific interactions among different ion channels, emergent properties can sometimes result in counter-intuitive observations that are perfectly explained by synergistic interactions. For instance, whereas an increase in sodium conductance is typically expected to enhance firing rate, it has been shown in certain populations that such increase results in decreased firing rate and gain. This has been attributed to synergistic interactions between sodium and delayed rectifier potassium conductances (Kispersky et al., 2012; Drion et al., 2015). Similarly, an increase in *A-*type K^+^ conductance is typically expected to reduce excitability, consequently reduce the amount of calcium entering into the cytosol and shift plasticity profiles to the right. However, under certain scenarios, owing to interactions of these conductances with the delayed rectifier potassium conductance, an increase in *A-*type K^+^ conductance results in enhanced calcium entering into the cytosol and leftward shifts to plasticity profiles (Narayanan and Johnston, 2010; Anirudhan and Narayanan, 2015). Similarly, in our case, we observed a counter-intuitive reduction in firing rate as a consequence of eliminating *A*- type K^+^ conductances. We argued that this was a consequence of enhanced activation of other potassium ion channels (KDR, SK and BK) by the larger and wider action potentials and by the consequently large influx of cytosolic calcium through voltage-gated calcium channels (Fig. 3).

Although the expression of *A-*type potassium condutcances in DG granule cells has been established through several lines of evidence (Beck et al., 1992; Sheng et al., 1992; Serodio and Rudy, 1998; Varga et al., 2000; Rhodes et al., 2004; Ruschenschmidt et al., 2006; Monaghan et al., 2008; Peng et al., 2013; Alfaro-Ruiz et al., 2019), heterogeneities in the impact of acutely eliminating *A-*type potassium channels have not been systematically assessed. Future studies should employ different techniques to acutely eliminate *A*-type potassium channels to assess potential heterogeneities of such elimination on neuronal firing rates. These experiments are especially important in light of the strong expression of calcium-activated potassium channels in DG granule cells, and their important roles in regulating intrinsic excitability of these neurons (Aradi and Holmes, 1999; Sailer et al., 2002; Brenner et al., 2005; Mateos-Aparicio et al., 2014). These acute blockade experiments should be coupled with systematic location-dependent recordings of *A*-type K^+^ currents (Hoffman et al., 1997) from the dendrites of DG granule cells, coupled with morphologically realistic computational models that account for such experimentally-assessed subcellular distribution of channels along the somatodendritic arbor (Beining et al., 2017). Such experiments would provide important insights about the specific roles of variable expression of ion channels, and unveil heterogeneities in terms how individual ion channels alter neuronal excitability as a consequence of spatiotemporal interactions with other ion channels along the somatodendritic arbor (Rathour and Narayanan, 2012b, 2014; Rathour et al., 2016).

From the standpoint of the impact of altered neuronal excitability to spatial encoding characteristics of DG neurons, it has been shown that hippocampal place maps are flexible, whereby the neural code of space remaps to mirror the animal’s behavioral experience (Dupret et al., 2010). One such remapping is where the firing rate of a place cell could undergo changes in response to environmental changes, with these changes also changing in a field-specific manner (Leutgeb et al., 2005; Leutgeb et al., 2007). Such rate remapping has been postulated to permit the distinctiveness of sensory events while maintaining the integrity of the spatial code (Renno-Costa et al., 2010). Our results show that modulation of intrinsic properties, either through neuromodulatory action or through activity-dependent plasticity could form a putative substrate for rate remapping in DG granule cells. Within this framework, field-specific rate remapping (Leutgeb et al., 2005; Leutgeb et al., 2007) could be achieved through differential neuromodulatory tones that are altered in a field-specific manner (as a potential consequence to specific behavioral associations to individual fields).

### Variability breeds robustness: Implications for the expression of local heterogeneities on ion channel regulation of response decorrelation

In our study, we demonstrated that knockout of individual ion channels alters channel decorrelation of the DG network, in a manner that is critically reliant on the specific heterogeneities expressed in the network. Although our analyses considered an extreme scenario where the ion channel was completely absent, the implications for changes to channel decorrelation as a consequence of modification to ion channel properties are several. Additionally, there are several detailed studies on the variable role of individual ion channels in altering single neuron physiology (see above), the extension of such analyses assessing the variable roles of individual ion channels (and consequent effects on intrinsic neuronal properties) to network scale functions have been far and few (Prinz et al., 2004; Morgan et al., 2007; Padmanabhan and Urban, 2010; Schneider et al., 2012; Padmanabhan and Urban, 2014; Yim et al., 2015). Our analyses, employing a systematic framework that accounts for several heterogeneities and multi-scale degeneracy in the emergence of single-neuron properties and network function (Mishra and Narayanan, 2019), highlights the importance of individual ion channels and their modulation to network-scale decorrelation *even* with the expression of the dominant afferent heterogeneities. Importantly, we demonstrate a critical role for local heterogeneities, especially the presence of immature neurons, in reducing the impact of ion channel knockouts on channel decorrelation.

The implications for these observations are three-fold. First, from a memory-encoding standpoint, it has been argued that DG neurons could act as engram cells, through changes in neuronal properties including cell-autonomous plasticity to membrane excitability (Zhang and Linden, 2003; Kim and Linden, 2007; Stegen et al., 2012; Yim et al., 2015; Gallistel, 2017; Titley et al., 2017; Josselyn and Frankland, 2018; Tonegawa et al., 2018; Rao-Ruiz et al., 2019). Neuron-specific changes in membrane excitability, potentially through alteration to ion-channel properties, during such plasticity would raise the question on how such changes in intrinsic properties would alter network-scale decorrelation properties. Our analyses suggest that the degree of decorrelation could alter as a consequence of such intrinsic plasticity, apart from providing a quantitative framework to address this question in a neuron-specific and channel-specific manner. Second, from a pathophysiological standpoint, acquired or inherited channelopathies are associated with several neurological disorders (Lehmann-Horn and Jurkat-Rott, 1999; Kullmann, 2002; Bernard et al., 2004; Hanna, 2006; Bernard et al., 2007; Beck and Yaari, 2008; Kullmann and Waxman, 2010; Terzic and Perez-Terzic, 2010; Poolos and Johnston, 2012; Brager and Johnston, 2014; Zhang et al., 2014; Johnston et al., 2016), and have been shown to be prevalent within the DG as well (Bender et al., 2003; Brenner et al., 2005; Stegen et al., 2009; Young et al., 2009; Surges et al., 2012; Kirchheim et al., 2013; Kohling and Wolfart, 2016). How does the network respond to such strong perturbations to ion channel composition, especially from the functional standpoint of response decorrelation? Our study suggests that such pathological changes in ion channel properties, and consequent changes to intrinsic excitability, would critically alter channel decorrelation in the DG. Our analyses also postulate that expression of local heterogeneities could help the network stay resilient to such large perturbations to ion channel densities.

Third, during the maturation process following generation of new neurons, it has been established that certain ion channels might not express during early stages of maturation (Ambrogini et al., 2004; Overstreet-Wadiche et al., 2006; Overstreet-Wadiche and Westbrook, 2006; Piatti et al., 2006; Lodge and Bischofberger, 2019). Therefore, the scenario analyzed here with the elimination of specific ion channels in immature neurons is equivalent to the absence of these channels during the maturation process. In all three cases, however, our analyses predict that the specific impact of these changes to channel decorrelation would depend on several factors, including the identity of the ion channel(s) involved, the answer to the question on what *other* component(s) changed, the nature of inputs to the network, the interactions of these altered components with other components in the network and the specific sets of heterogeneities expressed in the network.

### Limitations and future directions

A relatively small network (compared to (Mishra and Narayanan, 2019)) was employed in this study owing to the computational complexity of the knockout simulations, each involving different network configurations, simulations and analyses, spanning the 7 channel VKMs in GC and 2 channel VKMs in BC, and for the expression or absence of different forms of heterogeneities. Simulations in this study were performed and results analyzed for a total of 40 ((base model + 7 GC VKMs + 2 BC VKMs) × (4 sets of heterogeneity configurations)) distinct network configurations, with each configuration entailing a period of 1000 s virtual traversal (with a simulation integration time step *dt* of 25 µs), accompanied by correlation and firing rate analyses spanning these large time series arrays. However, it should be noted that the equivalence of results with small and large networks was established previously (Mishra and Narayanan, 2019).

In our study, we had analyzed the impact of complete elimination of individual ion channels. However, our conductance-based multi-scale modeling framework could be employed to assess the impact of perturbation to specific sets of channels that are observed under physiological or pathophysiological conditions. In such scenarios, the heterogeneous impact of changes could also be incorporated within the framework to make specific predictions on how network function would change under heterogeneous plasticity in different network components. Although our analyses here are with reference to channel decorrelation, future studies could explore the impact of individual ion channels on pattern decorrelation within the same framework, which would provide specific insights into the role of distinct forms of heterogeneities and different ion channels on pattern separation. Finally, the conclusions presented here could be electrophysiologically tested, both from the single-cell and network perspectives employing pharmacological agents to block specific ion channels or employing genetic methods to silence specific ion channels. In such analyses, interpretations should account for potential compensatory and activity-dependent plasticity mechanisms that follow the elimination of individual ion channels as well.

## Acknowledgments

This work was supported by the Wellcome Trust-DBT India Alliance (Senior fellowship to RN; IA/S/16/2/502727), Human Frontier Science Program (HFSP) Organization (RN), the Department of Biotechnology (RN), the Department of Science and Technology (RN), and the Ministry of Human Resource Development (RN & PM). The authors thank the members of the cellular neurophysiology laboratory for helpful discussions and for comments on a draft of this manuscript.

